# Complex compositional and cellular response of river sediment microbiomes to multiple anthropogenic stressors

**DOI:** 10.1101/2024.06.07.597903

**Authors:** Tom L. Stach, Aman Deep, Iris Madge Pimentel, Dominik Buchner, Mikayla A. Borton, André Soares, Jörn Starke, Till L.V. Bornemann, Philipp M. Rehsen, Jens Boenigk, Matthijs Vos, Florian Leese, Daniela Beisser, Alexander J. Probst

## Abstract

Rivers face constant anthropogenic stressors, resulting in significant changes in microbial community composition. What remains unclear is whether these changes render the microbiome better adapted to the stressed environment. Here, by subjecting 64 river-connected mesocosms to multiple stressors, we show that sediment microbiomes of small lowland rivers are highly sensitive to lowered flow velocity resulting in substantially altered community compositions not fully capable of compensating for the stressor effect within two weeks albeit having stable functions encoded in metagenomes. Transcriptomics revealed a systematic heat shock response in the community and a highly active, previously unknown anaerobic key stone species with great metabolic versatility. Increases in temperature (+3.5 °C) or salinity (+0.5 mS/cm) were outcompeted by lowered flow and elicited only minor responses at community or transcriptomic level with, *e.g.*, upregulation of the photosystem of chloroplasts. Following a two-week recovery period, transcriptomic stress responses vanished completely compared to control mesocosms, exemplifying the river microbiomes’ resilience. We conclude that, given the complex community responses at both the cellular and compositional level, maintaining natural river flow is vital to preventing energy loss and reduced microbiome activity in river sediments.

## Introduction

Stream ecosystems make up only 0.58 % of Earth’s non glaciated land surface area (Allen and Pavelsky, 2018), but are hotspots for biodiversity. While these ecosystems are particularly species rich, they are losing biodiversity at an accelerated pace globally, putting water security at risk (Vörösmarty et al., 2010). Although several large-scale frameworks such as the US Clean Water Act, the European Water Framework Directive, the UN’s sustainable development goals for 2030, and the Aichi Targets for 2020 are endorsed, the global situation has not substantially improved (Arthington, 2021; Tickner et al., 2020). Even positive developments seen at large scale in Europe for some indicators have come to a halt since 2010 (Haase et al., 2023). Thus, a key global task acknowledged in the new Kunming-Montreal Global Biodiversity Framework endorsed at 2022 United Nations Biodiversity Conference (COP 15) is to put freshwater ecosystems into conservation focus (Cooke et al., 2023). This recognition acknowledges the relevance of streams in providing essential services for nature including transport and cycling of carbon and nutrients on a longitudinal scale (Fasching et al., 2020; Tiegs et al., 2019) and pollutant removal (Hilderbrand et al., 2023). For humans, rivers provide numerous direct benefits, among them the provisioning of drinking water (Vermaat et al., 2016).

Microorganisms, *i.e.,* unicellular prokaryotic and eukaryotic organisms, play a crucial role in all of the above mentioned processes within streams. For example, Rodríguez-Ramos *et al*. revealed using genome-resolved metaproteogenomics complex riverine microbiomes being responsible for carbon and nitrogen cycling while impacted by viruses (Rodríguez-Ramos et al., 2022). Consequently, stream microbiomes were discussed as a central factor in climate change estimations as rivers were described as one of the “main sources of greenhouse gas (GHG) emission to the atmosphere” (Battin et al., 2023). For human needs, diverse microbiomes in various ecosystems can act as ‘shields’ against pathogens (Blaser and Falkow, 2009; Kennedy et al., 2002; Mahnert et al., 2019), but also reduce nutrient load and toxic substances before and during drinking water production from stream water (Shang et al., 2023; Sun et al., 2018).

A question that has so far been only poorly addressed is, how prokaryotic communities and their functions in stream ecosystems change in the Anthropocene under multiple stressors. As a stressor, we define any environmental variable that, due to anthropogenic interference, exceeds its normal range of variation with effects on organisms, communities and ecosystems (Townsend et al., 2008). The most prominent stressors to stream ecosystems are temperature increase, nutrient pollution, and hydromorphological changes (Nõges et al., 2016), whose impacts are particularly pronounced in densely urbanized areas (Tao et al., 2021). This situation is directly linked to microbial communities since experimental studies have identified direct positive links between their diversity and ecosystem functions (Biodiversity-Ecosystem Functioning (BEF) concept; Cardinale, 2011; Cardinale et al., 2012). While testing the effect of single stressors on microbial communities is straightforward and can be studied using lab experiments (Dang et al., 2021), field sampling (Havens et al., 2001), or meta-studies (Jurdzinski et al., 2023), disentangling the combined effects of multiple stressors is challenging due to the number of possible combinations and dependency of used parameters and models (Schäfer and Piggott, 2018) and is restricted to few studies (*e.g.,* Burdon et al., 2020; Romero et al., 2020). Thus, general principles of microbial community responses to multiple stressors and their recovery after stressor release remain scarce. However, this knowledge is crucial to identify combinations of stressors that are particularly harmful for microbial biodiversity and associated ecosystem functions due to potential synergistic interactions and thus crucial for remediation of stressed ecosystems (Brasseur et al., 2023; Hawley, 2018; Liu et al., 2020).

An experimental system to test multiple stressors in full-factorial settings with high replication while still under semi-natural conditions is the *ExStream* system, which has been widely used in stream ecology research (Beermann et al., 2018; Brasseur et al., 2023; Elbrecht et al., 2016; Nuy et al., 2018). For instance, Nuy *et al*. investigated the effect of multiple anthropogenic stressors on microbiota with a focus on the communities on leaf litter and biofilm tiles (Nuy et al., 2018). Their results already showed that stressors like salinization, fine sediment, and flow velocity have distinct response patterns for different microbes. Yet, they concluded that detailed taxonomic resolution and functional information were missing for better understanding the effect of stressors on complex river microbiomes (Nuy et al., 2018). In another mesocosm study, Romero et al. showed that low water level dominated the bacterial response and antagonistic effects to temperature and pesticides prevailed in the stressor phase (Romero et al., 2020). However, they determined that further studies are needed especially by moving from an indoor setup to field conditions. In general, the recovery of previously stressed microbiomes with a focus on the prokaryotic community in stream sediments has not been studied so far. A first synthesis of recovery patterns of disturbed microbiomes showed that aquatic communities tended to move away from their composition before disturbance, also emphasizing that recovery is environment-specific (Jurburg et al., 2024).

Taken together, the responses of microbial communities and their associated encoded and expressed ecosystem functions to multiple environmental stressors remain understudied. This is particularly true for riverine sediments. Given the essential role of the riverine sediment microbiome for key aquatic ecosystem functions, understanding responses to multiple stressors and their release is essential also for restoration and ecosystem management. Capitalizing on the BEF concept (Cardinale, 2011) and the Asymmetric Response Concept (Vos et al., 2023), a conceptual framework for the effects of multiple stressors on river biomes, we herein investigated river sediment mesocosms as a proxy for the hyporheic zone of rivers, which has been coined the “river’s liver” (Fischer et al., 2005). We applied three ubiquitous anthropogenic stressors, *i.e.,* increased temperature and salinity, and lowered flow velocity, in a full-factorial design using the *ExStream* system at the Boye stream in North Rhine-Westphalia, Germany. We investigated the sediments during the stressor application phase, and additionally elucidated the recovery of the microbiome after stressor release. Combining this setup with 16S rRNA gene amplicon analysis, genome-resolved metagenomics, and metatranscriptomics, we aim at (i) identifying compositional and transcriptional community responses, (ii) revealing sentinel organisms and keystone organisms under multiple stressor conditions, and (iii) inferring functional gain or loss due to stressor increase. Our results demonstrate a complex interplay of anthropogenic stressors on river microbiome composition and functional response with the major result that compositional microbiome changes in river sediments cannot compensate for the stressed ecosystem necessitating regulatory mechanisms including heat shock responses in bacteria.

## Results

For investigating the individual and combined effects of three different anthropogenic stressors on river microbiomes, a full-factorial mesocosm approach was applied (***Figure 1a-c***). After an acclimation phase of 20 days of the mesocosm system, stressors were applied for 14 days followed by a recovery phase without stressors for another 14 days. The applied stressors included temperature increase (+3.5 °C; denoted as “T+” vs. “T0”), salinity increase (+0.5 mS/cm; “S+” vs. “S0”), and flow velocity change (3.5 vs. 14.2 cm/s; “V-“ vs. “V0”; ***Figure 1d***). Samples for 16S rRNA gene amplicon, metagenomic, and metatranscriptomic analyses were taken after the stressor and recovery phase, respectively.

**Figure 1:**
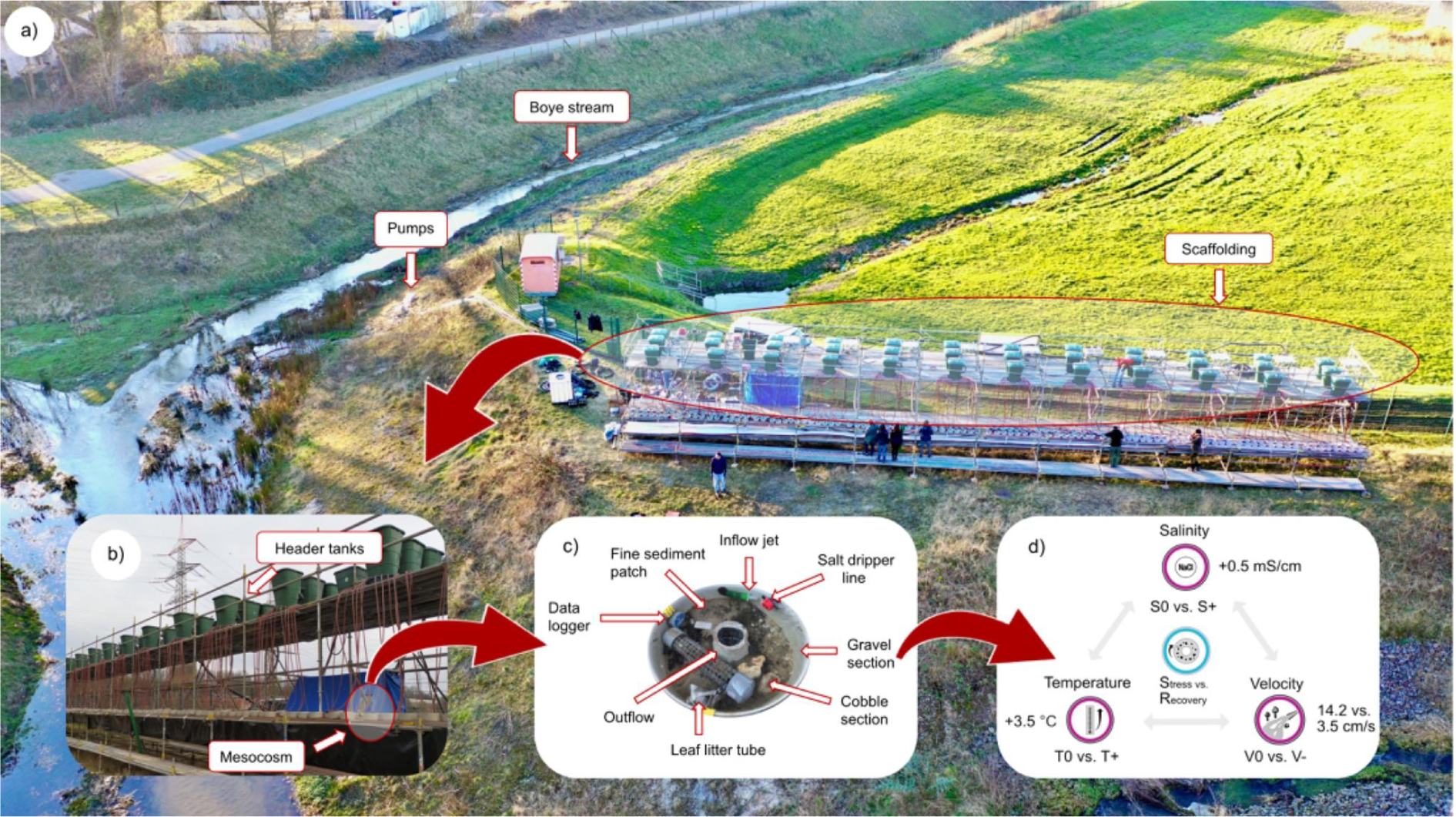
Experimental design of the highly parallelized mesocosm setup called ExStream. (a,b) Stream water was directly pumped from the Boye stream to the scaffolding with overall three ExStream setups, of which one with 64 mesocosms was used for this study. (c) Arrangement of the circular mesocosms used in this ExStream experiment. During the stressor phase, the low flow velocity was achieved by removing the white cap at the inflow jet. Salinity was added by a salt dripper line, and temperature was increased by mixing in-flowing river water with heated river water (constantly measured by data logger). Sediment sampling was done in the fine sediment patch. (d) Stressor combination applied in full-factorial design with a minimum of three replicates for each phase respectively. Image provided by Philipp M. Rehsen.

### Lowered flow velocity as a significant driver of sediment microbiome composition outcompeting effects of temperature and salinity increase

Extracted DNA from sediment was used for marker gene analysis based on 16S rRNA gene amplicons and normalized *ribosomal protein S3* (*rpS3*) gene sequences. Multivariate statistics revealed a highly significant microbial community shift due to flow velocity change for both 16S rRNA gene and *rpS3* gene analysis after stressor application (adonis2; p=0.001, Tab. 1). Flow velocity change also resulted in the highest chance-corrected within-group agreement (*i.e.*, the greatest difference between community compositions) amongst all tested stressors in metagenomic data (MRPP; A=0.008; p=0.001). However, combinations of stressors, *e.g.*, temperature combined with salinity and lowered flow velocity, revealed different community effects than certain stressors alone, *e.g.*, lowered flow velocity.

**Table 1:**
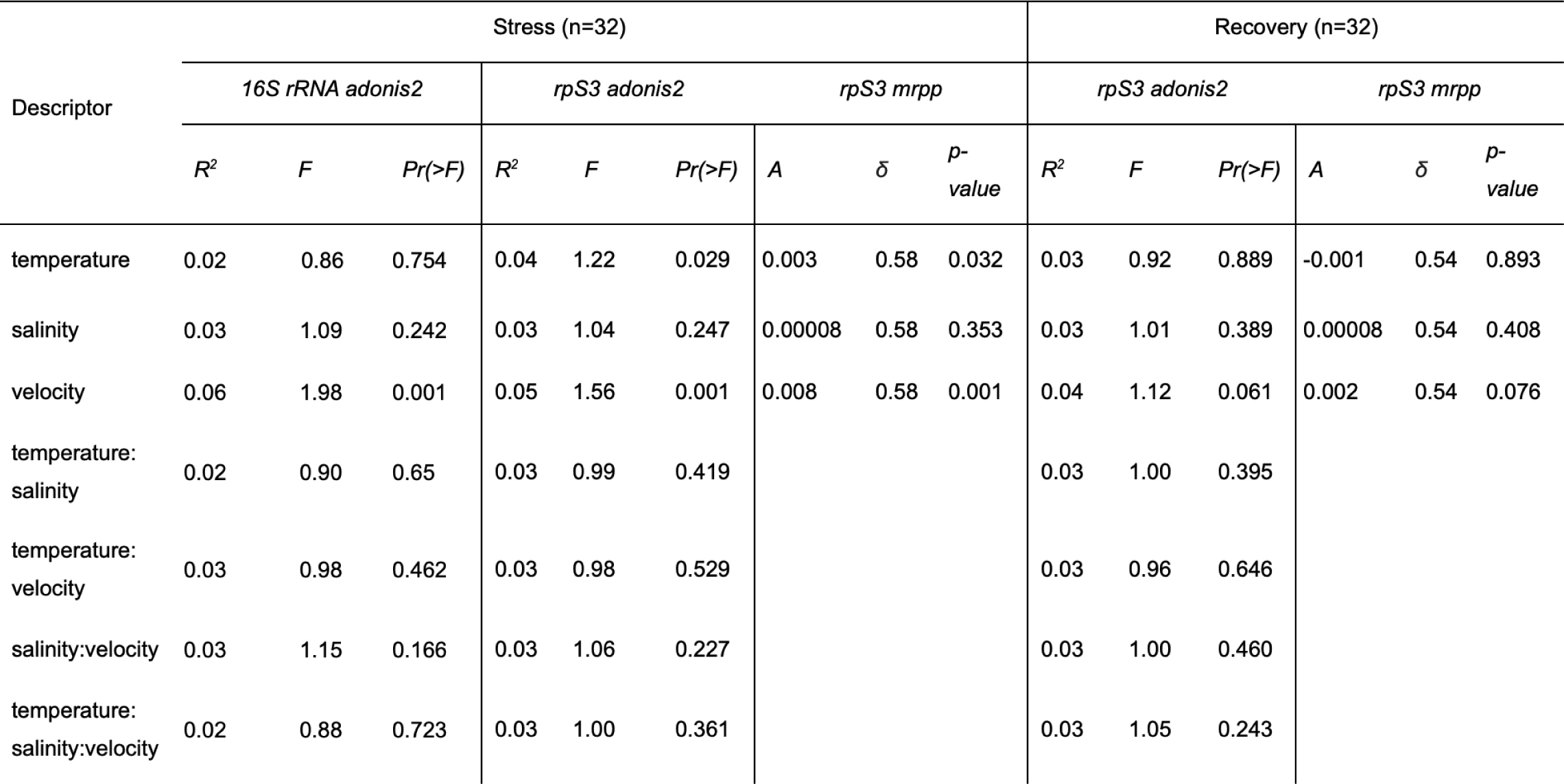
Statistical testing of stressor impact on microbial community structure based on relative abundance of representative 16S rRNA gene amplicon data and representative rpS3 gene sequences from metagenomic sequencing. Adonis2 was run with 999 permutations and a model including all single factors and interaction terms based on the marginal effects of the terms as test design. MRPP was grouped for descriptors individually and ran with 999 permutations. DF=1 for descriptors.

Prokaryotic communities exposed to lower flow velocity clustered closely together when visualized with Non-Metric Multidimensional Scaling (NMDS) based on 16S rRNA gene data (***Figure S1***). This was accompanied with a uniform increase of diversity compared to all other mesocosm setups, including both controls and other stressor combinations (Shannon: p=0.0009, Simpson: p=0.008; ***Figure S2***). Amplicon-based microbiome analysis is well known to be affected by primer-bias and PCR-bias which may lead to an unsuitable measure for accurate river bacteriome analyses (Stach et al., 2023). Microbiome analysis based on *rpS3* gene sequences extracted from de novo assembled metagenomes aims at overcoming these issues by being PCR-free, and covering eukaryotic gene signatures and a higher diversity of uncultured prokaryotic microorganisms (Sharon et al., 2015). Consequently, *rpS3* gene analysis also uncovered temperature increase as an additional stressor with a significant microbiome change (adonis2; p=0.029 in *rpS3* data versus; p=0.754 for 16S rRNA gene data).

Recovery of the microbiome structure after flow velocity stress was well documented by non-significant community diversity statistics (Tab. 1) as well as by NMDS based on *rpS3* gene data (***Figure 2***). Whereas channels treated with low flow velocity group closely together after the stressor phase (***Figure 2a***), this pattern disappeared after recovery (***Figure 2b***).

**Figure 2:**
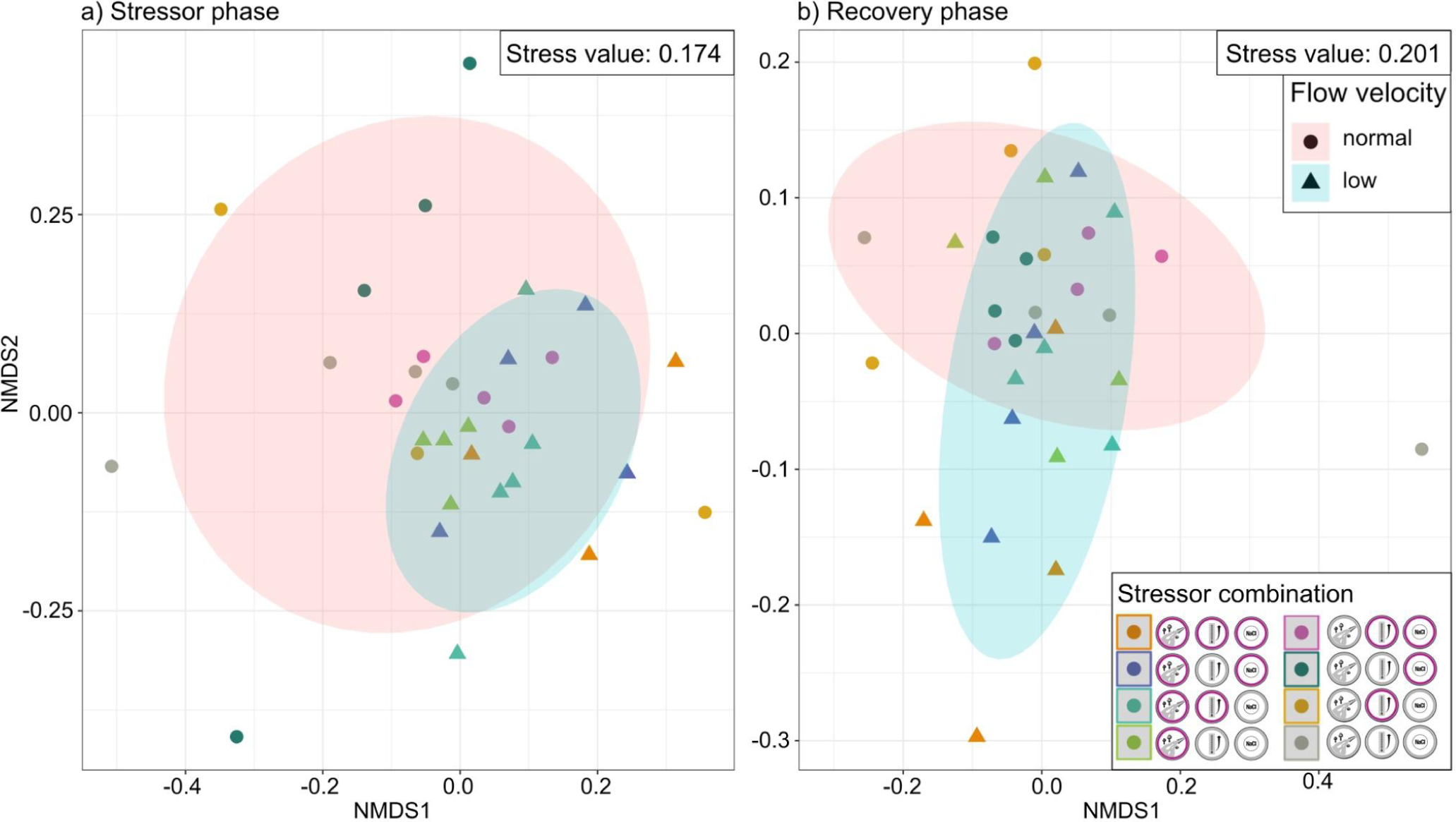
Non-metric multidimensional scaling (NMDS) of normalized abundances of representative rpS3 gene sequences from metagenomic sequencing (Bray-Curtis dissimilarity matrix after rarefaction (100 iterations)) separated for stressor (n=32) and recovery phase (n=32), respectively. Stressor combinations were grouped by velocity treatment (ggplot2, ellipse level=0.75) indicating a unifying effect of low flow velocity to the stressed microbiome.

To disentangle the relationship of stressor treatments from one another, the mean pairwise Bray-Curtis dissimilarity between two sets of treatments against all other stressor sets was tested (***Figure 3***) demonstrating significant differences in stressed mesocosms compared to the control. Regarding the recovery phase, no significant differences were detected when comparing its control with previously stressed channels highlighting the validity of the mesocosm approach presented herein by being able to track the community development into a new steady state.

**Figure 3:**
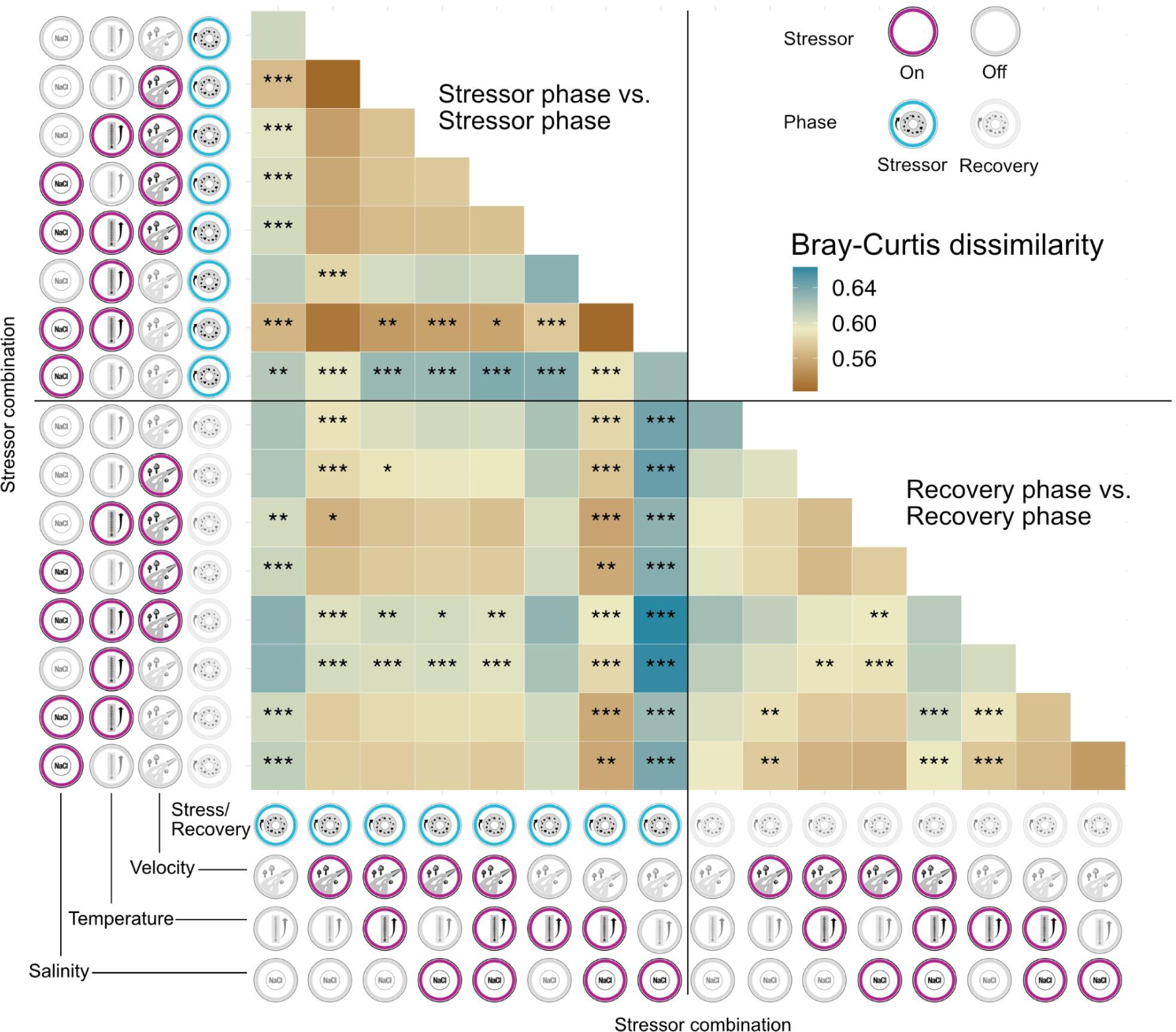
Heatmap displaying the pair-wise mean differences in Bray-Curtis distances compared to all stressor sets (n=64, all at least in triplicates) based on rarefied relative abundance of representative rpS3 gene sequences. For each comparison, the dissimilarity values of respective stressor sets compared to all other stressors were extracted (Wilcoxon test with Bonferroni correction for multiple-testing, asterisks indicate significance level). For example, the control of the stressor phase and channels treated with low flow velocity showed a highly significant difference when the mean of all dissimilarity values of these channels were compared.

All treatments with low flow velocity had a comparably low mean dissimilarity to each other (0.52-0.58) and were not significantly different compared to other treatments, when testing for individual stressors alone. Thus, all river mesocosms with low flow velocity treatment developed similar microbiomes. Interestingly, salinity as a single stressor resulted in not only high dissimilarities compared to other stressors but also within its own replicates (0.63), and significantly different relationships to all other treatments. Unlike flow velocity, salinity did not cause a uniform microbiome response explaining its insignificant differences compared to other stressor sets in PERMANOVA testing. When combining salinity increase with temperature increase, mean dissimilarities, however, dropped and were comparable to treatments with flow velocity change. After recovery, pronounced patterns as seen during the stressor phase drastically dropped with 17 treatment comparisons (47 %) showing significant differences after stressor impact compared to nine after recovery (25 %) indicating a high microbiome resilience at the taxon level.

### Indicator organisms for specific stressors in the river sediments include highly abundant microbes

The stream sediment microbiomes harbored a high biodiversity, and this was also reflected on the read level by Nonpareil3 analyses with a mean Nonpareil diversity (Nd) value of 23.32 ± 0.13 (n=64) (Nonpareil3 curves are displayed in Figure S3) of which sequencing covered between 45.04 % and 63.16 % (average of 54.7 ± 3.77 %). This indicates that our approach only covered about half of the sequence diversity in our samples, likely attributed to incomplete eukaryotic genomes in our samples that are generally known to be of high complexity compared to prokaryotic genomes. This limitation was overcome by assembling metagenomes of all 64 mesocosms and creating an *rpS3* gene sequence database comprising 14,255 lineages, which were also used for multivariate analyses (see above).

To understand the detailed stress responses at organism level, we first investigated the top ten organisms based on *rpS3* gene sequence abundances focusing on the differences between stressed and recovered mesocosms (***Figure 4***). Overall, Betaproteobacteria, Gammaproteobacteria, and eukaryotic Bacillariophytha dominated the microbiomes across all treatments, yet multiple organisms with identical taxonomic annotation were present among the top ten organisms due to our clustering approach geared towards resolving microdiversity rather than species (clusters based on 99% gene similarity). Nine out of the ten most abundant *rpS3* gene clusters showed a significant increase in relative abundance during recovery compared to stressor application providing further confirmation for the asymmetric response of river microbiomes to stressors resulting in different microbiomes after the recovery phase. The relative abundance of top abundant representative *rpS3* gene sequences in control channels fell in most cases below the mean of all samples (***Figure 4***). We conclude that the top most abundant taxa, which demonstrated greater tolerance to the applied stressors based on relative abundance measures, leveraged this advantage during the recovery phase. They probably outcompeted the more sensitive taxa and derived greater benefit from the removal of stressors.

**Figure 4:**
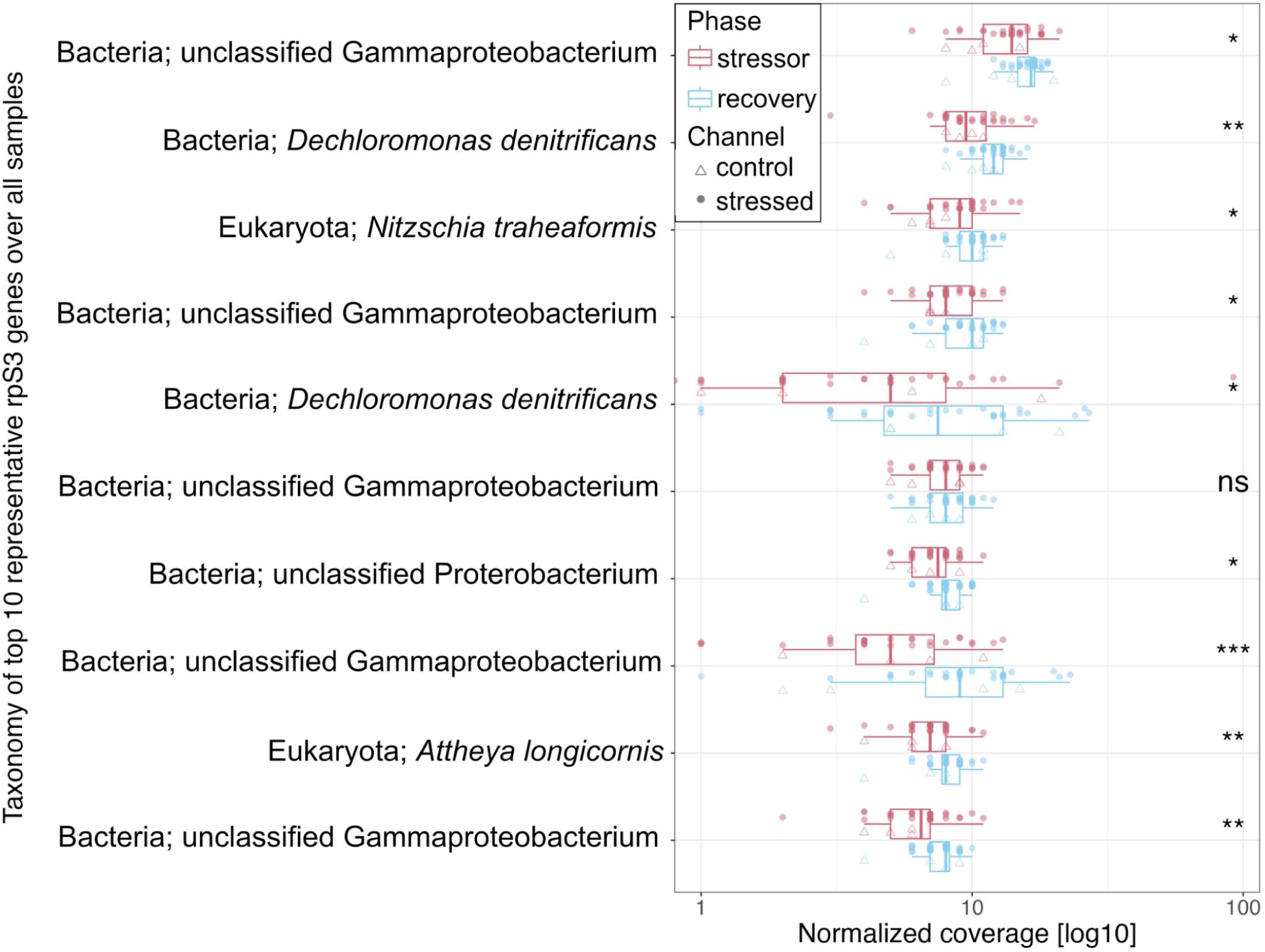
Box-Whisker plots of top ten most abundant microbes across all samples (ordered from top to bottom based on abundance) with sequencing-depth normalized relative abundances of representative rpS3 gene sequences in stressed and recovered mesocosms with controls labeled as triangles. Nine out of ten rpS3 gene clusters showed significant increases in relative abundance when comparing stressor and recovery phase (Wilcoxon test). Figure with full taxonomy is available as Figure S4.

To disentangle microbiome response at taxon level, *i.e.*, the relative abundance change of *rpS3* gene signatures depending on stressor application, we identified overall 110 potential indicator organisms (TukeyHSD post-hoc testing; adj. p-value < 0.01; ***Figure 5***, Figure S5, and Figure S6). Altogether, 62 organisms across 9 phyla responded either negatively or positively to decreased flow velocity, 19 of which increased or decreased only if at least one additional stressor was applied (***Figure 5***).

**Figure 5:**
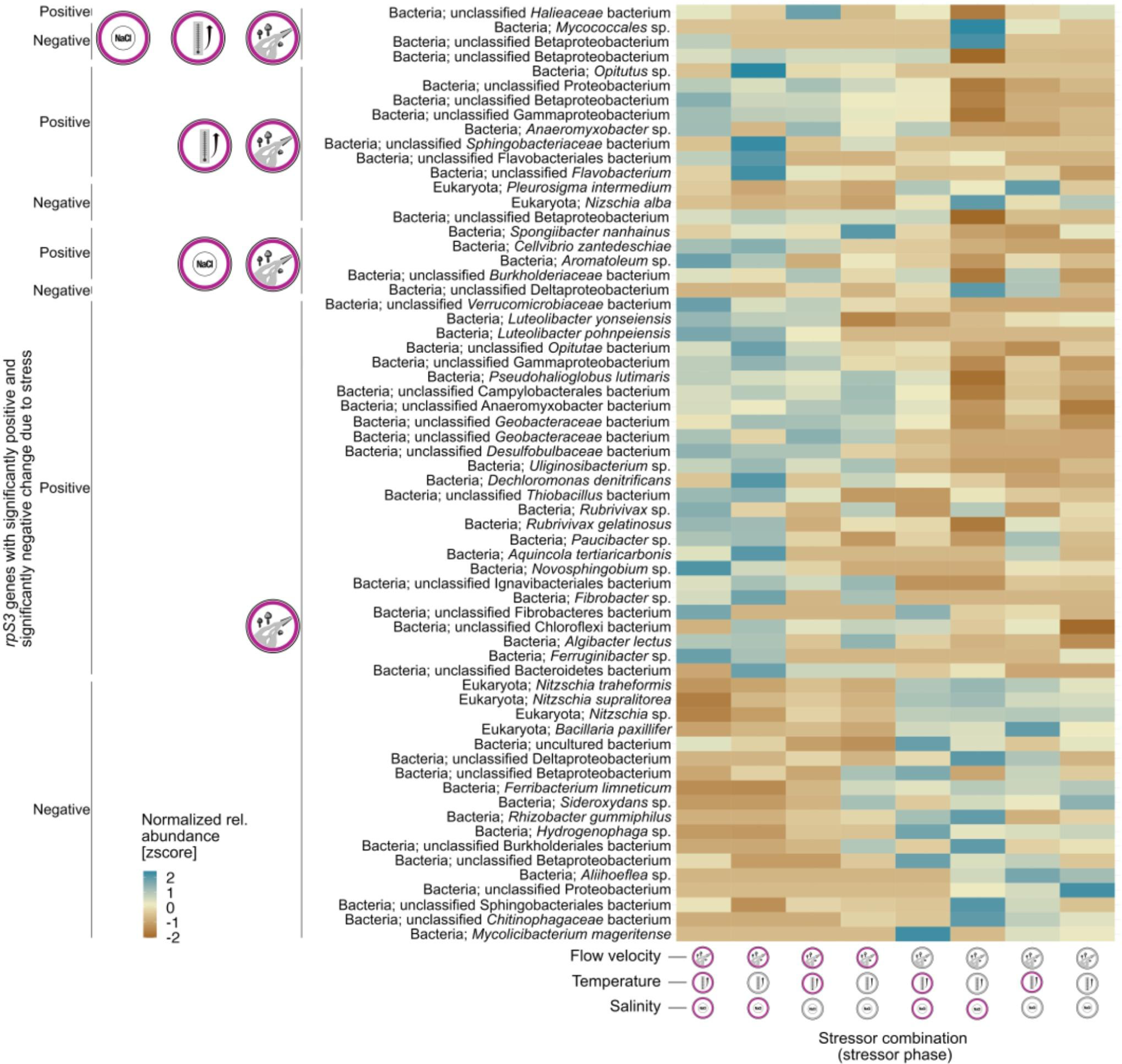
Microbial taxa sensitive to flow velocity change based on sequencing-depth normalized representative rpS3 gene sequences (ANOVA followed by TukeyHSD; adjusted p-value < 0.01). If multiple stressor effects per taxon were significant, only the lowest p-value was chosen. For example, the taxon at the very bottom, Mycolicibacterium mageritense, was negatively affected by low flow velocity, although its individual relative abundance was not uniform across stressor sets. For relative abundance change of organisms due to salinity or temperature change, please see Figure S5 and S6, respectively.

One main indicator taxon for slow flow velocity are diatoms, *e.g., Nitzschia* sp., which responded negatively to velocity decrease, whereas the bacterium *Dechloromonas denitrificans* responded positively, each of them being part of the top ten organisms in the sampled river sediments (***Figure 4***). In case of temperature increase (Figure S6), certain thermophilic bacteria like *Schlegelella thermodepolymerans* responded positively as well as three representative sequences annotated as *Luteimonas* spp.. Parasitic organisms like Candidatus Gracillibacteria or CPR SR1 bacteria, as well as an eukaryotic endosymbiont of *Kryptoperidnium foliaceum* responded negatively to temperature increase indicating that they or their hosts are either outcompeted by others or did not perform well after a 3.5 °C temperature increase. Salinity increase affected the microbiome to a smaller degree compared to the other stressors (Figure S5), which is in agreement with multivariate analyses (Table 1). Yet, one proteobacterial lineage typically found in marine environments responded positively to salt increase solely, *i.e., Haliea* sp., or in combination with temperature increase and velocity reduction, *i.e., Halieaceae*. The results of putative thermophiles responding to temperature increase and salt-adapted organisms responding to salt change links physiological knowledge to changes in gene signatures and confirms our analysis approach. The 110 identified indicator organisms expand our knowledge of microbial indicators for stressed river ecosystems substantially.

### Organism-specific functional changes are related to stressor application and recovery

To link biodiversity change to ecosystem functions (BEF), genome-resolved metagenomics was coupled to metatranscriptomics, enabling the identification of differentially expressed genes of metagenome-assembled genomes (MAGs) due to stressor application. MAGs with a minimum completeness of 75 % and maximum contamination of 10 % from all metagenomic samples and the public GROWdb (which is based on river MAGs to increase number of potential hits in mapping) were dereplicated resulting in a set of 63 at least medium-quality MAGs according to MIMAG standards (Bowers et al., 2017). The main share of MAGs were annotated as Proteobacteria. Across all MAGs, a high strain heterogeneity (46.5 % ± 30.40 %) was detected (***Figure 6a***), which is similar to certain soil microbiomes (Banchi et al., 2023; Ma et al., 2023; Viacava et al., 2022) and explains the generally low genome recovery of MAGs from river sediments. When testing for response to applied stressors based on read mapping of metagenomic data, three MAGs were significantly positively impacted by low flow velocity (Tukey-HSD; max. p.-adj. < 0.01) and one MAG by temperature increase (Tukey-HSD; p.-adj. < 0.001), being consistent with signatures found for 16S rRNA gene amplicon and *rpS3* gene sequence data (Tab. 1). The latter belonging to the class of Bacteroidia was only present in samples with increased temperature (Figure S7). This sentinel species encoded seven genes that were significantly differentially expressed due to temperature increase (see Figure S8). Three of those genes encoded for uncharacterized proteins, three other for T9SS type A sorting domain-containing protein, and the last one for a HU family DNA-binding protein. The latter two encode for crucial functions that are important for the integrity of the cell and metabolism functioning. For instance, the T9SS type proteins are known to be a factor in virulence (Barbier et al., 2020), motility (Nakane et al., 2013), and S-layer formation (Tomek et al., 2014) of Bacteroidetes. HU DNA-binding proteins on the other hand are essential for DNA coiling, repair, and transcription (Berger et al., 2010).

**Figure 6:**
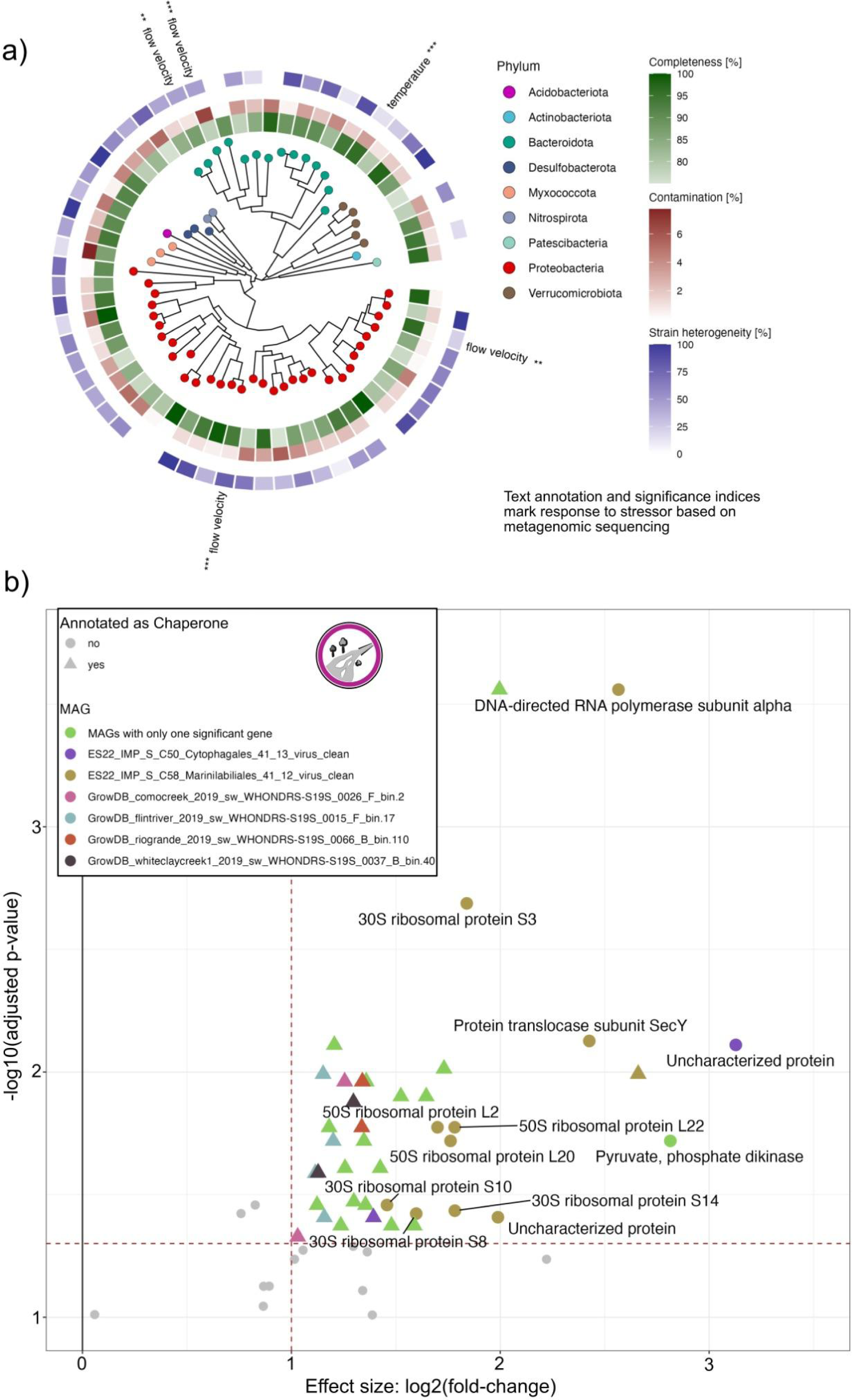
Phylogenetic, stressor response, and transcriptomic analysis of MAGs from ExStream experiment combined with MAGs from GrowDB (M. Borton et al., 2022). a) Dereplicated MAGs (n=63; completeness over 75 % and contamination under 10 %) were dominated by Proteobacteria. When tested for different abundances due to applied stressors (n=32 channels), four MAGs significantly positively responded to lowered flow velocity and one to increased temperature (text annotation with significance index as asterisks). b) Metatranscriptomic reads were mapped to the set of MAGs and tested for differential expression due to lowered flow velocity using DESeq2 (n=32 for stressor phase; log2(fold-change) > 1 and adjusted p-value < 0.05; Love et al., 2014). 17 MAGs encoded for only one gene responding to flow velocity, respectively. Chaperones were identified as indicator genes for lowered flow velocity. For stressor combinations after the stressor phase and after the recovery phase in general, no differentially expressed gene could be identified. Differential expression due to temperature increase is shown in Figure S8.

In case of low flow velocity, 41 genes present in at least one of the MAGs were significantly upregulated (***Figure 6b***). From these genes, 28 encoded for chaperone-related proteins and seven for ribosomal proteins. One MAG which was significantly affected on metagenome level by low flow velocity, annotated as *Burkholderiacea,* even responded with two differentially expressed chaperone genes. These results demonstrate that several organisms responded with upregulating protein production (translation via ribosomes for which ribosomal proteins are necessary) but also increased their stress response in the form of chaperones which are necessary for correct protein folding (Hartl et al., 2011; Tokuriki and Tawfik, 2009).

The MAG with most differentially expressed genes (in short, C58_Marinilabiliales_41_12) belonged to the family of *Prolixibacteraceae*. Also, the expression of its *rpS3* gene was significantly upregulated at low flow velocity according to metatranscriptomic data, which did not show an increase based on metagenomic data and was only detected in three samples based on metagenomic read mapping (***Figure 5***). This indicates low abundance but high activity of this organism rendering it a typical keystone species. We investigated the encoded metabolism of the keystone species’ genome (84.8% complete, 2.17% contamination, ∼4 Mbps) using manual annotation in the MicroScope platform (Vallenet et al., 2009) deciphering the potential carbon, nitrogen, sulfur and phosphorus sources of the organism (***Figure 7***). The genome encoded a variety of transporters for the uptake of cations, ammonia, sulfate, phosphate, amino acids, acetate, and sugars pinpointing at a heterotrophic organism, which was further corroborated by the presence of a near-complete glycolysis and TCA cycle for synthesis of molecular building blocks for amino acids and breaking down organic carbon to carbon dioxide. Interestingly, the conversion of acetate to acetyl-CoA, which is essential for processes such as the complete fatty acid biosynthesis encoded in the genome, could theoretically occur through acetate uptake and conversion via acetylphosphate or through de novo synthesis via gluconeogenesis. The major sulfur source for amino acids appeared to be assimilatory sulfate reduction, while nitrogen sources were either ammonia uptake via transporters or strictly anaerobic nitrogen fixation via NifHD (three subunits encoded next to each other and next to a nitrogenase iron protein). Regarding energy conservation, NAD(P)H can be produced from either oxidation of organic compounds via TCA cycle or via hydrogen oxidation (hydrogenase Hnd with subunits A-D). We also identified potential electron acceptors based on gene annotations. While the membrane-bound nitrate reduction to ammonia via nitrite is fully encoded in the genome for energy conservation, we also identified tetrathionate to thiosulfate conversion with the latter potentially being reduced to sulfite (thiosulfate/3-mercaptopyruvate sulfurtransferase potentially contributing its product to assimilatory sulfate reduction, see above) or sulfide (via a thiosulfate reductase with a membrane-bound cytochrome subunit). Subunits of an encoded NADH oxidoreductase likely energize the membrane via oxidation of NADH. Additional energization of the membrane could be achieved via an encoded Rnf-complex, although we could not identify a reaction for reduction of ferredoxin encoded in the genome. Given that the MAG is not complete, we could either miss the respective gene(s) or not be able to annotate them. However, electron bifurcation along with the encoded hydrogenase might also be a source for reduced ferredoxin. Overall, this heterotrophic and chemolithotrophic keystone organism showed a great versatility of encoded pathways in the genome being able to harness a variety of different nutrients and energy sources, which potentially relates to its high activity. The lowered flow velocity as a stressor likely limited oxygen supply into the sediment resulting in microhabitats that enabled this strictly anaerobic microbe to thrive.

**Figure 7:**
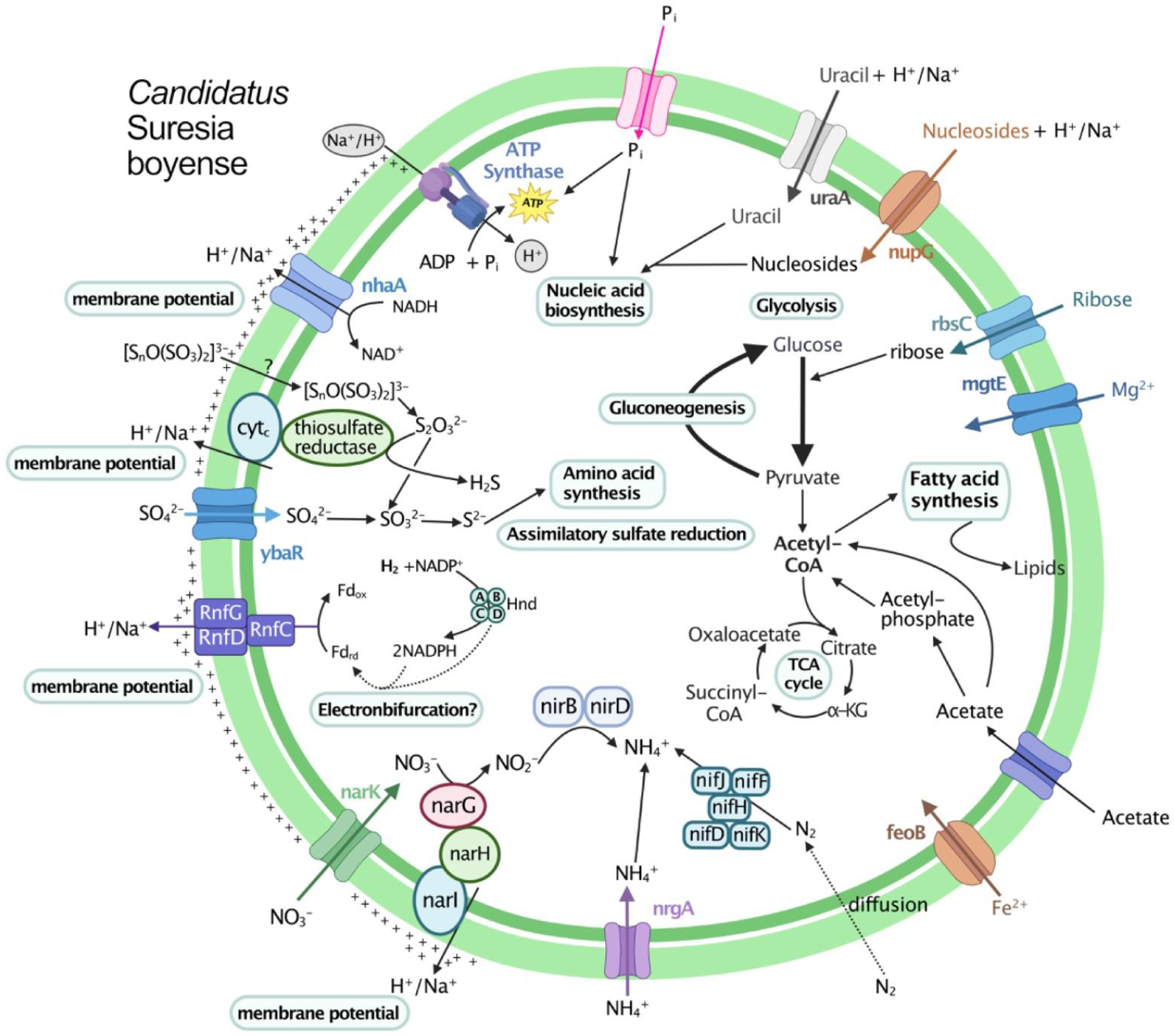
Metabolic capacity of an identified key organism in the analyzed microbial community of the Boye: MAG C58_Marinilabiliales_41_12 (GTDB taxonomy; phylum Bacteroidota, family Prolixibacteraceae) named Candidatus “Suresia boyense” after Bernd Sures who pioneered using the Boye stream as a model ecosystem for restoration. We used one MAG for manual reconstruction of this metabolic map (n=1) in the MicroScope platform (Vallenet et al., 2009) followed by visualization created with BioRender.com. For details on metabolism, please see the main text.

### Encoded ecosystem functions remain stable at larger scale but are sensitive at gene expression level

Given that genome-resolved metagenomics does not capture all functional capacities of microbiomes, we also analyzed the metagenomic and metatranscriptomic data at the gene level. No significant changes due to stressor application could be identified using multivariate statistics based on normalized relative abundance of metabolic modules present in metabolic assemblies as detected by METABOLIC (Zhou et al., 2022), which included carbon and sulfur cycling, nitrogen turnover, and oxygen metabolism. Since absence of significant difference does not imply significant similarity, relative abundance of encoded functions were tested for equivalence using the TOSTER package (Lakens, 2017) (Figure S9) revealing that the amount of stressors applied had an impact on the detected number of equally shared functions. Specifically, when all stressors were applied, no functions were significantly equivalently encoded compared to the control or channels with only one stressor. When temperature was increased as a sole stressor, functions were only equally shared with channels that also had temperature as a stressor. Given the absence of significant differences of prokaryotic functions and the frequent detection of their similarity, we conclude that these were mostly stable across all stressor tests. However, Eukaryotic community changes, *e.g.,* the decrease of diatoms during reduced flow velocity (see above), imply that at least primary production of the system was substantially affected.

Focusing on the expression pattern of genes of prokaryotic origin (West et al., 2018) across all metagenomes (***Figure 8a-f***), we identified the salinity-driven upregulation of 99 genes of which 51 were annotated as “photosystem” of plastids. Only 33 of genes were affected by other stressors, including temperature increase (12), low flow velocity and salinity increase in combination (12), or all stressors combined (9). Across all metatranscriptomes, gene expression was most affected by the stressor flow velocity resulting in 221 genes (twelve downregulated, 209 upregulated), of which 38.5% had an annotation string containing “chaperon”. Five upregulated genes were related to heat-shock proteins (“hsp20”, “Small heat shock protein”, and “heat shock protein”). Comparing the response patterns of stressor combinations with each other based on functional groups (***Figure 8a***) low flow velocity activated typical stress response mechanisms of microorganisms, and salinity increase also led to a distinct response. Other stressors, alone or in combination, did not trigger specific responses, if having an impact at all. Thus, specific responses due to lowered flow velocity and increased salinity were gone when combined with other stressors. While prokaryotic functions remained stable at large scale, applied stressors activated responses at gene expression level. Since no differentially expressed genes were detected after the recovery phase, we conclude that the gene expression-based stress response existed only for a very short period of time until the stress was released.

**Figure 8:**
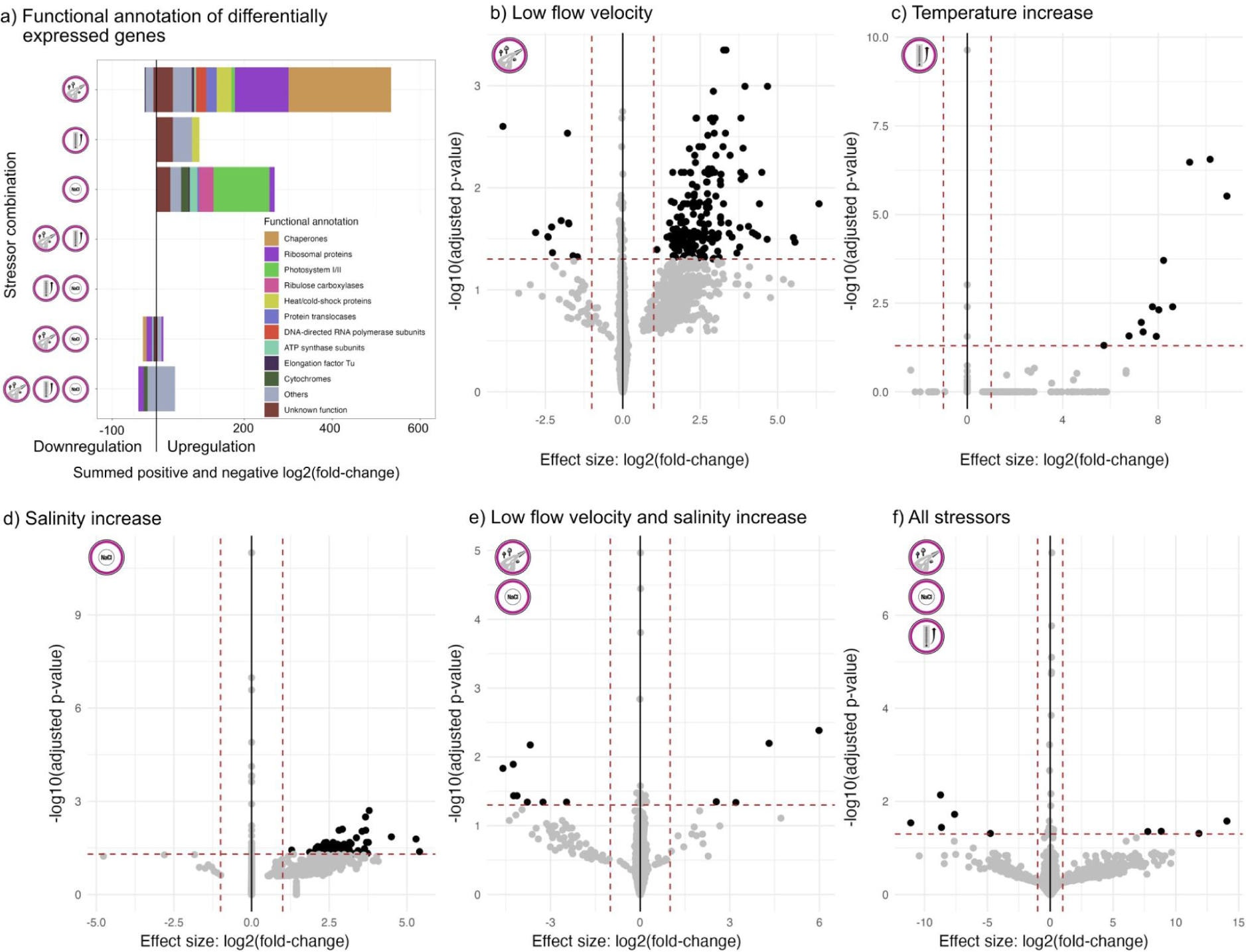
Functional metatranscriptomic data. a) Overview of differentially expressed genes and their functional annotation after stressor application. Up- and downregulated genes were summed up respectively and manually grouped into functional groups revealing that low flow velocity and salinity increase lead to specific responses while other stressors are rather nonspecific if having a response at all. b-f) Metatranscriptomic reads were mapped to all prokaryotic genes clustered at 90% similarity and tested for significance (dashed lines represent log2(fold-change) > 1 and adjusted p-value < 0.05). Plots refer to the treatments after the stressor phase and only those with significantly different gene abundances are shown. Lowered flow velocity results in drastic upregulation of prokaryotic genes (209; 38.5 % annotated as chaperones). Salinity increase results in upregulation of photosystem genes encoded on plastids. There were no significantly differentially regulated genes after the recovery phase.

## Discussion

Stream ecosystems are globally under stress (Dudgeon et al., 2006; Harding et al., 1998). Efforts to restore stressed streams often fail in returning them to a natural state due to a lack of systematic understanding how previous stressor exposure shapes recovery trajectories after stressor release (Vos et al., 2023). Capitalizing on the biodiversity and ecosystem function (BEF) concept (Cardinale, 2011) and our previously published asymmetric response concept (ARC) (Vos et al., 2023), we tested these theoretical frameworks specifically for river sediment microbiomes. The ARC suggests that main mechanisms in stream community assembly, *i.e.,* stress tolerance, dispersal, and biotic interactions, play roles at differently pronounced levels during stressor increase and after stressor release. We show that integration of stress tolerance of microbiomes is key for predicting their response to, *e.g.*, lowered flow velocity. Investigation of most abundant taxa across channels suggests that stress tolerance during the stressor phase led to having benefits after stressors were removed with a headstart compared to more sensitive taxa considering relative abundance measures. In general, lowered flow velocity caused canalisation (reduced variability) while salinity as a stressor enhanced noise (increased variability) in the microbial community that however got canalized when high temperature was added as a second stressor (Berlow, 1997). The response to salinity was only detectable at taxa and expression level without a uniform response in community change (Adonis test) as observed for velocity and temperature, which can also be explained by the history of our model stream being previously impacted by mining and used as an open sewer system potentially resulting in a microbiome adapted to higher levels of salt concentration (Winking et al., 2014, 2016). Similar findings have been identified for eukaryotic communities and ecosystem functions in a preceding experiment (Madge Pimentel et al., 2024). This would suggest a long-term adaptation of river microbial communities, which should be tested for in the future.

Across our full-factorial outdoor mesocosm study, bacterial river communities from sediments responded significantly to lowered flow velocity not only by a change in community but also by upregulating chaperones, which is a clear sign for energy-demanding stress responses (Fourie and Wilson, 2020; Zahrl et al., 2007). Such processes can consequently drain cellular energy (Goloubinoff et al., 2018; McCarty et al., 1995) and likely reduce the microbiomes’ fitness and capacity for nutrient turnover in rivers. The upregulation of photosystem encoding genes due to salinity increase, similarly found along a river-to-sea continuum (Tee et al., 2021), pinpoints to an increase of energy needed to cope with the energy stress that the microbial community is faced with. Taken together, these adaptations have implications for the primary production in streams, not only due to the discussed gene expression shifts, but also due to abundance shifts, *e.g,*. a lower abundance of diatoms due to lowered flow velocity. Given the fact that rivers and streams are substantially involved in carbon dioxide burial from terrestrial input (0.3 Pg C yr^-1^; Battin et al., 2023) and diatoms account for approximately 20% of total primary production on Earth (Field et al., 1998; Tréguer et al., 2018), we posit that decreased flow velocity of rivers may negatively impact primary production which is, however, a vital ecosystem service in the context of climate change (river gross primary production 0.07 Pg C yr^-1^; Battin et al., 2023). Thus, allowing a river to have natural flow variations, with both fast and slow flowing sections, could greatly improve provision of essential ecosystem functions (Hilderbrand et al., 2023; Huang et al., 2021) and recovery from stressed phases.

In summary, our results show that flow velocity (and to a lesser extent also temperature) significantly change the stream sediment microbiome at community and organism level. While there is redundancy of encoded functions, the microbiome was forced to acclimatize at least short-term through transcriptomic stress response, especially to lowered flow velocity and increased salinity. We did not detect a community fully functional without stress response, so the tremendous bacterial diversity available through dispersal did not deliver a set of new dominant species that could handle the experimental conditions as a stress-free environment in the tested time frame of two weeks. Consequently, the restructured microbiome after two weeks of stressor application of lowered flow velocity was not perfectly adapted to the new stress conditions and still needed to upregulate its heat stress response in the bacterial population. We conclude that such a heat stress response of bacteria is a substantial energy sink draining energy from important ecosystem processes like nitrogen turnover rendering the populations less productive. However, these changes were reverted after two weeks of recovery highlighting the resilience of the river sediment microbiome in general. Comprehensively, this study identified one major requirement for restoration procedures of sediment microbiome functioning, *i.e.*, allowing rivers to flow freely within natural ranges of velocity, so that these communities can perform their respective ecosystem services, and a healthy river microbiome is maintained.

## Materials and Methods

### Study site

The outdoor mesocosm system *ExStream* (***Figure 1***) was set up at the Boye stream (51.5533 °N, 6.9485 °E) and operated from March 4, 2022 to April 21, 2022. The Boye stream, part of the Emscher River system in North Rhine-Westphalia, Germany, had a history of anthropogenic stressors stemming from coal mining, industrial wastewater, and open sewage discharge from the early 19th until 2019 and is now fully restored since 2021 (Perini and Sabbion, 2017; Winking et al., 2014).

### Experimental system and mesocosm setup

The *ExStream* system is an open mesocosm system where a continuous supply of water from the Boye stream flows into multiple circular channels (Madge Pimentel et al., 2024; Piggott et al., 2015). Briefly, this setup consisted of a two-level scaffold with sediment trap tanks and header tanks on the top that supply the mesocosms on the bottom level (***Figure 1b***). Stream water was first pumped to the sediment trap tanks to reduce the velocity of the water and settle the sediment down to the bottom. Consequently, overflowing water from the sediment trap tanks entered one of four header tanks with 16 connected mesocosms each. Water supply from the header tanks to each channel was given by gravitational force. All pipes connected with mesocosms had a shut-off valve to adjust discharge to 2.1 L/min. The mesocosms had a diameter of 25 cm and a volume of 3.5 liters. The *ExStream* system is an open system, *i.e.*, after passing through the experimental system the water is directed back into the stream (see Madge Pimentel et al. 2024).

In every mesocosm **(*Figure 1c***), 1 liter of sediment from Boye stream (0-1 mm) was added, along with 100 ml slurry of fine particulate organic matter (from a small tributary stream close to the Boye stream; 51.5627885 °N, 6.9150225 °E), serving as an initial carbon source in order to represent the typical substrate of the studied river (Madge Pimentel et al., 2024). Gravel (quartz stones, 6-8 mm, store-bought) was positioned behind the inflow jet with three pebbles (quarry stones, 40-80mm, store-bought) on one side of the gravel section and another pebble (quarry stones, 32-56 mm, store-bought) on the other side. Organic matter was added to the mesocosms by using air-dried alder leaves (*Alnus glutinosa*) collected from the coordinates 51°20′59.09″N, 7°10′14.03″E. Water temperature was constantly measured in 8 representative channels using data loggers, conductivity and water temperature was measured separately using external probes. The leaves were packed into fine mesh bags before being added to the mesocosms. A check for potential prokaryotic contamination of the mesocosms based on full-length 16S rRNA gene Nanopore sequencing was done beforehand showing no detectable contamination (see Supplementary Information).

### Experiment design

The experiment was divided into three phases, acclimatization, stressor, and recovery phase. During the acclimatization phase of 20 days, all 64 mesocosms were operated without manipulation. Subsequently, during the stressor phase, stressors including temperature, salinity, and velocity were applied to mesocosms for 14 days in a full-factorial design with eight replicates per treatment combination. Specifically, the temperature was increased by +3.5 °C (denoted as “T+” vs. “T0”) compared to ambient stream temperature. Salinity was increased by 0.5 mS/cm (“S+” vs. “S0”) using NaCl input via dripper lines, and velocity was lowered from 14.2 to 3.5 cm/s (“V0” vs. “V-”) by removing the in-flow jet. The first endpoint sampling of 32 mesocosms was done after the 14-day stressor phase. Four mesocosms per treatment were removed and used for analysis. The remaining 32 mesocosms continued to run without stressor application for 14 days in the subsequent recovery phase. Thus, each treatment for the stressor and recovery phase respectively had four replicates. Due to an oversight during initial setup, the number of replicates differed for four stressor combinations., *i.e.*, N=3 for V-S+T+ (velocity lowered, salinity increased, temperature increased) and V0S0T+, and N=5 for V-S0T+ and V0S+T+ during the stressor phase. Unpredictable, yet natural, fluctuations in streamflow and water level occurred between days 7 and 10 of the stressor phase. These fluctuations, caused by heavy rainfall in the catchment area, resulted in temporary pump blockages. Consequently, stressors were only applied for a limited period (12 hours) on day 8. Normal conditions resumed on the morning of day 10, allowing for the continuation of the stressor phase as planned.

### Temperature as stressor

Temperature increase in mesocosm was introduced via heated water from an electric mobile heating system (triMobil EHZ36, maximum capacity: 36 kW) using filtered (125 μm stainless steel filter that was automatically flushed every 10 min for 15 s) stream water as input. The heated water was drained into 10 L buckets installed above the header tank. Heated water was mixed with cold water from the stream to achieve +3.45 °C (treatment means ± SD: T_ambient_ = 8.71 ± 0.06 °C, T_warming_ = 12.16 ± 0.08 °C, n = 4) degree water in the mesocosm via a static mixer (Madge Pimentel et al., 2023). Header tanks that received only unheated stream water were instead supplied with an amount of filtered stream water that was equivalent to the heated water.

### Salinity as stressor

Salt tablets (Claramat, > 99.9 % NaCl) mixed with stream water were supplied through dosage pumps to mesocosm via dripper lines to receive an increase by 0.529 mS/cm (treatment means ± SD: EC_ambient_ = 0.842 ± 0.006 mS/cm, EC_salt_ = 1.343 ± 0.151 mS/cm, n=32), corresponding to 154.1 mg/L added chloride. The salinity setup closely resembled the methodology outlined in the Madge Pimentel et al. (2024).

### Velocity as stressor

The velocity of water was lowered by removing the inflow jet (Madge Pimentel *et al*., 2024). The lowered flow mesocosms had 3.50 ± 3.32 cm/s (n = 4, measured in the middle of the water column, in the first quarter of a mesocosm) and the normal flow velocity with an inflow jet mounted had 14.25 ± 7.59 cm/s (n = 4). The normal flow velocity corresponded to a typical flow velocity of the studied stream type, *i.e.*, a small sand-dominated lowland river (type 14 according to Dahm et al. (2014)). Consequently, the lowered flow velocity was designed to be outside of the normal flow velocity of the model river type.

### Sampling procedures and sample processing

At the endpoint of the stressor and recovery phases, sediment from the respective 32 mesocosms were collected. Sediment cores were taken using a sterile, cut 50-ml syringe capturing the whole depth of the sediment patch right in front of the inflow jet (***Figure 1c***). Subsequently, the samples were manually homogenized, aliquoted, shock-frozen in liquid nitrogen, and stored at −80 °C until further processing.

### DNA and RNA extraction and sequencing

DNA was extracted for metagenomic and 16S rRNA gene amplicon sequencing from 0.5 g sediment input. For DNA extraction, samples were mixed with 0.1 and 0.5-mm diameter glass beads and 100 μl Proteinase K, 5 μl RNAse A, and 900 μl TNES (for buffer and reagents see materials in Buchner (2022e). Then, samples were bead-beaten for 2 min at 2400 rpm in a Mini-Bead-Beater 96 (Biospec Products, Bartlesville, USA). Samples were incubated at 56 °C and bead beaten at 1400 rpm for 20 minutes. Lysates were divided for replicates and DNA was extracted following the spin column protocol using a vacuum manifold described in Buchner (2022b). DNA clean-up was performed with carboxylated-magnetic beads and PEG-NaCl buffer following the protocol described in Buchner (2022c) with 40 μl DNA input and 80 μl of clean-up solution. Eluted DNA was split and used for metagenomic sequencing directly or library preparation for 16S rRNA gene amplicon sequencing. RNA was extracted from sediment for metatranscriptomic sequencing using phenol/chloroform/isoamyl alcohol as described in Buchner (2022a). Sequencing for metagenomes, metatranscriptomes, and 16S rRNA gene amplicons was performed at CeGat Gmbh (Tübingen, Germany).

### 16S rRNA gene amplicon sequencing and analysis

For 16S rRNA gene analysis, DNA was amplified with a two-step PCR approach. 10 μl reaction volume was used per sample for the first PCR with Multiplex PCR Plus Kit (Qiagen) with primers 515f/806r (Apprill, A *et al*., 2015), and 1 μl of DNA input. After the cleanup of the first PCR product (Buchner, 2022c), 2 μl of DNA was used for the second PCR. Cycling conditions for the first and second PCR are given in Supplementary Table S3. Bead-based normalization protocol was used for DNA normalization which resulted in a 2 ng/μl concentration (Buchner, 2022d). The pooled libraries were concentrated using a spin-column clean-up protocol with a final volume of 100 µL (Buchner, 2022b). PCR replicates were produced from each sample and these libraries were then subjected to paired-end sequencing (2 x 250 bp) on Illumina NovaSeq (CeGat Gmbh, Tübingen).

Raw reads were processed with the Natrix2 workflow (Deep et al., 2023). In brief, the pipeline included primer removal, assembly with Pandaseq (v2.11, Masella et al., 2012) and filtering the paired-end reads with a alignment threshold score of 0.9 and sequence length with a minimum of 100 bp and a maximum of 600 bp. Dereplication (100% sequence similarity) and removal of chimeric sequences was done with cd-hit (v 4.8.1, Fu et al., 2012); erroneous sequences were removed with a split sample approach using AmpliconDuo (v1.1, Lange et al., 2015). Resulting sequences were clustered with Swarm (v2.2.2, Mahé et al., 2014) into OTUs which were aligned against the Silva database (v138.1, Quast et al., 2012) by using Mothur (v1.40.5, Schloss et al., 2009) for taxonomic classification. MUMU (https://github.com/frederic-mahe/mumu), an update for LULU (Frøslev et al., 2017) was used for post-clustering. For quality assessment, PCR replicates of each sample were consolidated by eliminating OTUs exclusive to one of the replicates and combining the reads from the remaining replicates. Additionally, number of reads for OTUs identified in negative controls were subtracted from all individual samples (46 OTUs). The read counts were normalized with the variance stabilizing transformation (VST) function from DESeq2 package (Love et al., 2014). These transformed read counts were then converted into PhyloSeq (McMurdie and Holmes, 2013) objects with metadata and taxonomic information for further statistical testing and plotting with ggpubr (Kassambara, 2023), tidyverse (Wickham et al., 2019), and vegan (Oksanen et al., 2013).

### Metagenomic sequencing, quality control, assembly, and annotation

Metagenomic sequencing was performed using the Illumina DNA PCR-Free Prep protocol on a NovaSeq 6000 with a minimum sequencing depth of 30 Gbp (150 bp paired-end reads; CeGat Gmbh, Tübingen). Metagenomic reads were quality checked and trimmed using BBduk (Bushnell, https://jgi.doe.gov/data-and-tools/bbtools/bb-tools-user-guide/) and Sickle (quality score ≥ 20 and minimum read length ≥ 20 bp) (Joshi and Fass, 2011). The covered microbial diversity by metagenomic sequences was estimated using Nonpareil3 (v3.4.1; kmer mode) (Rodriguez-R et al., 2018).

Metagenomic paired-end reads were assembled per sample using MEGAHIT v.1.2.9 (Li et al., 2016) with preset ‘meta-large’. For resulting contigs at least 1 kbp in length, open reading frames were predicted using Prodigal (parameters: -m -p meta; Hyatt et al., 2010) and annotated using DIAMOND blast (DIAMOND version 2.0.15; blastp --fast -e 0.00001 -k 1) (Buchfink et al., 2021) against FunTaxDB (version from Aug. 2022; Bornemann et al., 2022), which is based on UniRef100 (Suzek et al., 2007). Taxonomy was assigned at the contig level based on all proteins detected. Specifically the lowest taxonomic rank was assigned when more than 50% of the protein taxonomies agreed (Bornemann et al., 2022). The average contig coverage was calculated based on mapping quality-checked reads to the assembly using Bowtie2 in sensitive mode (Langmead and Salzberg, 2012). For all contigs, length and GC content were calculated.

### Ribosomal protein S3 marker gene analysis

Based on a phylosift Hidden Markov Model (HMM) set (DNGNGWU00028; date 20.01.2022) (Darling et al., 2014) *rpS3* gene sequences were identified in metagenome assemblies (hmmsearch (v3.2), e-value-cutoff of 1E-28). Additionally, annotation results using FunTaxDB (see described above) were searched for *rpS3* genes, and the respective taxonomy was attached. Positive hits for *rpS3* genes from both approaches were searched against a *rpS3* set for archaea and bacteria from GTDB r207 (Parks et al., 2022) using USEARCH (Edgar, 2010) reporting the best hit with an e-value of 1E-5 or better (or reporting unclassified if no hit with these criteria was found).

For statistical analyses of *rpS3* gene sequences across samples, *rpS3* gene sequences were clustered using USEARCH (-cluster_fast -id 0.99). Per cluster, the centroid of the cluster was chosen as the representative sequence if flanked by at least 1 kb in both directions on its contig. Otherwise, a non-centroid longest sequence that was respectively flanked could be extended or just the longest sequence was selected. The contigs were then trimmed to the respective length including the flanking regions in order to improve mapping results (Figueroa-Gonzalez et al., 2020). Then, quality-filtered reads were mapped with bowtie2 (Langmead and Salzberg, 2012) representative *rpS3* gene sequences excluding reads shorter than 100 bp (reduction of 3.2% ± 0.34% on sequence length) to ensure a sensitive and specific mapping. Mappings were filtered for three mismatches or less to exclude random alignments and coverage was used for statistical analysis.

To test for the significance of stressors on microbial communities, coverage values were rarefied to the lowest summed coverage of all samples (ES22_IMP_S_C04, sequencing depth: 26.1 GB; summed coverage 2051) with 100 iterations and permanova test (adonis2) was performed based on Bray-Curtis dissimilarity with 999 iterations (used R packages included vegan (Oksanen et al., 2013), permute (Simpson, 2022) and GUniFrac (Chen et al., 2023)). When running the rarefaction for the stressor phase individually, the lowest summed coverage differed to channel ES22_IMP_S_C38 (sequencing depth: 29.8 GB; summed coverage 2272). The mean dissimilarity between all treatments was calculated, tested using the wilcox test, and resulting p-values were adjusted using the Bonferroni method. Based on the dissimilarity matrix, nonmetric multidimensional scaling (NMDS) with grouping per stressor at a level of 0.75 and Multi-Response Permutation Procedure (MRPP) tests were performed using the vegan package in R (Oksanen et al., 2013).

Additionally, each representative sequence was tested for stressor effects after the stressor phase only. For that, mapping-based relative abundances were normalized based on the lowest sequencing depth, and each sequence was tested as follows. A two-way anova was performed and the resulting p-value was adjusted for multiple testing using the Bonferroni method. Subsequently, for representative sequences with a p-value lower than 0.05, a post-hoc Tukey-HSD test was run.

### Binning of assembled metagenomes and curation of MAGs

Binning contigs into MAGs followed the approach described in Stach et al., 2023. In brief, MetaBAT2 (Kang et al., 2019), ABAWACA (Brown et al., 2016), and MaxBin2 (Wu et al., 2016) were used as binning tools. For the latter, a cross-mapping over all samples was done using Bowtie2 (Langmead and Salzberg, 2012), and both marker sets were used. In the end, an optimized set of bins per sample was produced using DASTool (Sieber et al., 2018), and resulting MAGs were manually curated using uBin v0.9.20 (Bornemann et al., 2023).

To ensure that no non-prophage viral sequences were included in the curated MAGs, viral sequences in all contigs were identified using VirSorter2 (v2.2.3; --high-confidence-only; Guo et al., 2021), deepvirfinder (v1.0.0; −l 1000; Ren et al., 2020), and VIBRANT (v1.2.1; Kieft et al., 2020). Viral sequences classified non-prophage with minimum 25 % completeness by checkV (Nayfach et al., 2020) were removed from the MAGs.

### Incorporation of publicly available MAGs from other river ecosystems

Already binned MAGs from an external database, *i.e.,* GROWdb (M. Borton et al., 2022; M. A. Borton et al., 2023), were included in the following analyses. The aim was to reduce the chance of missing MAGs due to assembly or binning problems originating from the high complexity and diversity of metagenomic samples.

Viral-clean MAGs from all 64 mesocosm metagenomes and MAGs received from GROWdb were dereplicated using dRep v3.4.3 (Olm et al., 2017). For that, Checkm2 (Chklovski et al., 2023) was run for all MAGs and the output was used for genome statistic calculation by dRep with a minimum completeness of 75%, maximum contamination of 10% and identity of secondary clusters of 95%. Coverage for dereplicated MAGs was calculated using the “quick-profile” module of inStrain v1.7.6. (Olm et al., 2021) using Bowtie2 mappings as input. For the following analyses, MAGs were counted as present with a breadth greater than 0.5, with breadth being defined as the fraction of the genome covered by at least one read. Coverage data for MAGs was used to identify MAGs responsiveness to stressor effects during stressor phase as described for *rpS3* gene sequences above. Taxonomic annotation and an unrooted tree were inferred using GTDB-Tk (v2.1.0, r207) workflow *de_novo* and taxonomic classifications were verified via the *classify_wf* mode (Chaumeil et al., 2020). Plotting of phylogenetic tree was performed with the help of ggplot2 (Wickham, 2016), ggdendro (Vries and Ripley, 2022), ggtree (Yu, 2020), ggtreeExtra (Xu et al., 2021), rcartocolor (Nowosad, 2018) and patchwork (Pedersen, 2023). Functional annotation of the final MAG set was performed using DRAM (v1.4.6; (Shaffer et al., 2020) and final quality assessment of MAGs according to Bowers et al., 2017 was done as described in (Stach et al., 2023). A detailed annotation for our keystone MAG (C58_Marinilabiliales_41_12) was done in MicroScope (Vallenet et al., 2009).

### Functional annotation and equivalence testing

Metabolic pathways were annotated to the metagenomic assemblies based on METABOLIC (v4.0) (Zhou et al., 2022) using the METABOLIC-G.pl mode. Relative abundance of HMM-modules was calculated by the mean coverage of contigs which encoded the respective genes and normalized on sequencing depth. Response to stressors was tested using multivariate statistics including adonis2 (Oksanen et al., 2013) and stressor combinations were tested for equivalent abundance of functional modules using the TOSTER package in R (Lakens, 2017). Taking the given number of mesocosms per treatment, the higher and lower equivalence bounds were calculated to achieve 33% power as suggested previously (Simonsohn, 2015).

### Metatranscriptomics RNA sequencing and analysis

Total RNA extracts were sequenced using the Illumina TruSeq Stranded Total RNA with Ribo-Zero PLUS Kit for rRNA depletion. Raw metatranscriptomic sequences were processed with a custom snakemake-based workflow (Köster and Rahmann 2012) (https://github.com/adeep619/Vasuki). In this workflow, raw sequences were quality filtered using cutadapt (v3.2, Martin, 2011) with phred score 20 and minimum length 50 bp. Ribosomal RNA sequences were removed from the quality checked and adapter-trimmed sequences using Ribodetector (Deng et al., 2022). Filtered mRNA sequences were mapped to genes from dereplicated MAGs and clustered genes. For the latter, all contigs from all metagenomic assemblies were filtered for prokaryotic origin using EukRep (v0.6.6, West et al., 2018) and genes were predicted using prodigal in meta mode. Subsequently, resulting genes were clustered using MMseqs2 (v1806c0c8f7aa4365f9f72c8ea51e947d1e93ccd9, lineclust) at 90% sequence identity (Steinegger and Söding, 2017). Reads were mapped to representative gene sequences using Bowtie2 (--sensitive) (Langmead and Salzberg, 2012), and counts were calculated using CoverM (Aroney et al., 2024). Differential expression analysis was performed in R using the DESeq2 package (Love et al., 2014) using all stressors and interactions in the test design (smallestGroupSize = 3, minimum five channels over smallestGroupSize, FDR/alpha=0.05). Shrunken log2 fold changes were calculated with type “apeglm” (Zhu et al., 2019), and taxonomic or functional annotations were added to those genes with thresholds of: alpha < 0.05 and abs(log2FoldChange) > 1. Differentially expressed genes were grouped into functional groups manually, if a functional group was present less than five times respective genes were called “Others”.

## Acknowledgments

This study was performed in context of the Collaborative Research Center (CRC) RESIST and analyses were mainly done in Projects A01 and A04, funded by the German Research Foundation (DFG) CRC 1439/1; project number 426547801. Tom L. Stach was supported by the German Academic Scholarship Foundation. We acknowledge support from the Open Access Publication Fund of the University of Duisburg-Essen and Project Deal (Elsevier and MPDL Services GGmbH, 2023). The authors would like to thank all involved student helpers, doctoral students, and supporting staff for their contribution to *ExStream*. Ken Dreger is acknowledged for exemplary server administration.

## Author contributions

The project was conceived and supervised by A.J.P. and Da.Be.. T.L.S. performed metagenomic and metatranscriptomic analysis after read polishing. A.D. performed 16S rRNA gene analysis and metatranscriptomic analysis until read polishing. Both contributed equally to sampling, data visualization, and writing of the manuscript. The mesocosm experiment was conceptualized and led by I.M.P., P.M.R., and F.L. Extraction and processing of nucleic acids was done by Do.Bu. External metagenomic data was derived from the Genome Resolved Open Watersheds (GROW) database, an effort supported by M.A.B. Sampling was supported by J.S.. A.S., T.L.V.B., and J.S. supported bioinformatic processing, data visualization, and data analysis. All authors revised the final version of the manuscript.

## Supplementary results

### Prokaryotic community analysis based on 16S rRNA gene amplicon sequencing data

**Figure S1:**
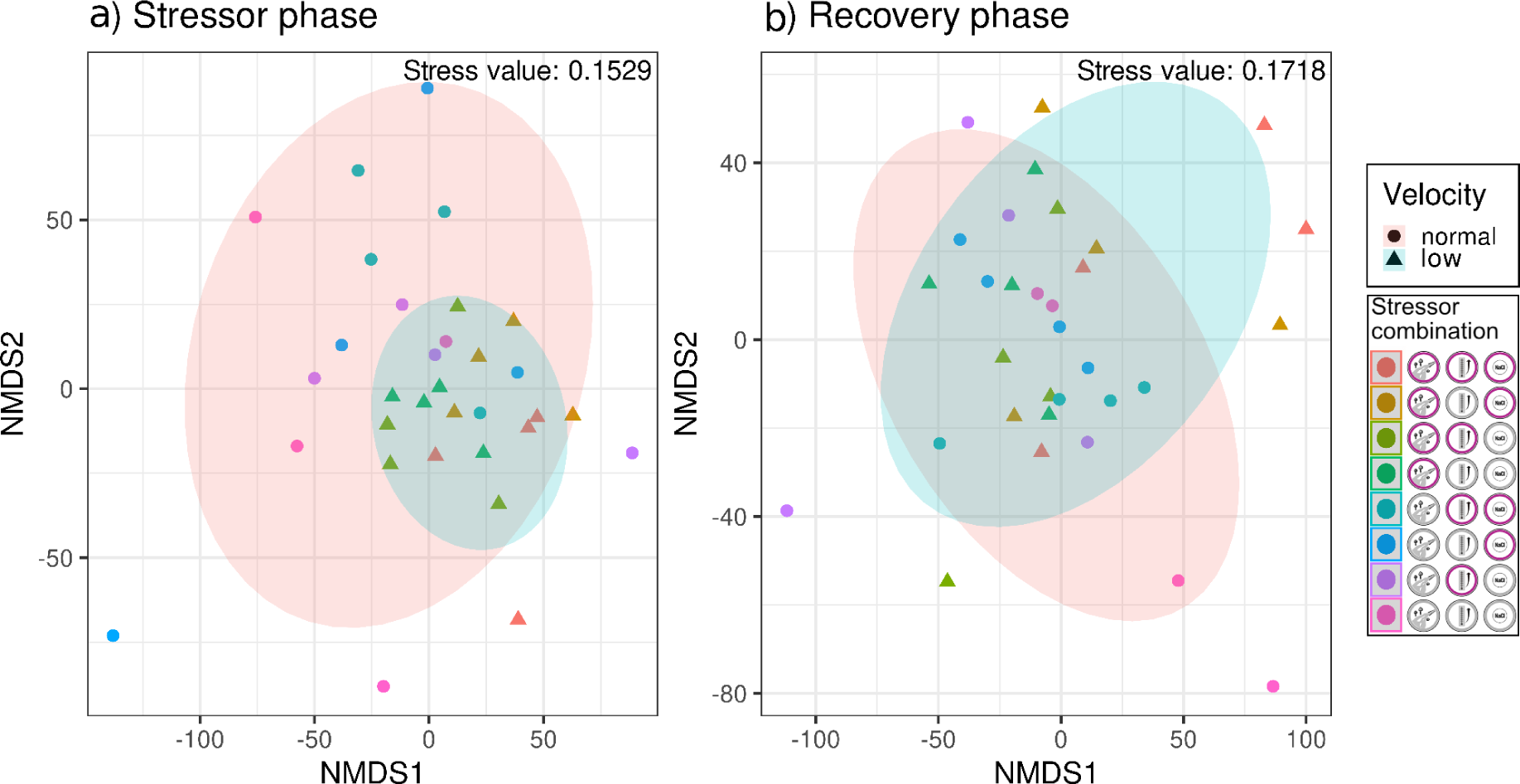
Non-metric multidimensional scaling (NMDS) of OTUs from 16S rRNA amplicon sequencing (Bray-Curtis dissimilarity matrix) separated for stressor (n=32) and recovery phase (n=32), respectively. Stressor combinations were grouped by velocity treatment (ggplot2, ellipse level=0.75) indicating a unifying effect of low flow velocity to the stressed microbiome.

### Significance testing of 16S rRNA gene data after recovery phase

**Table S1:**
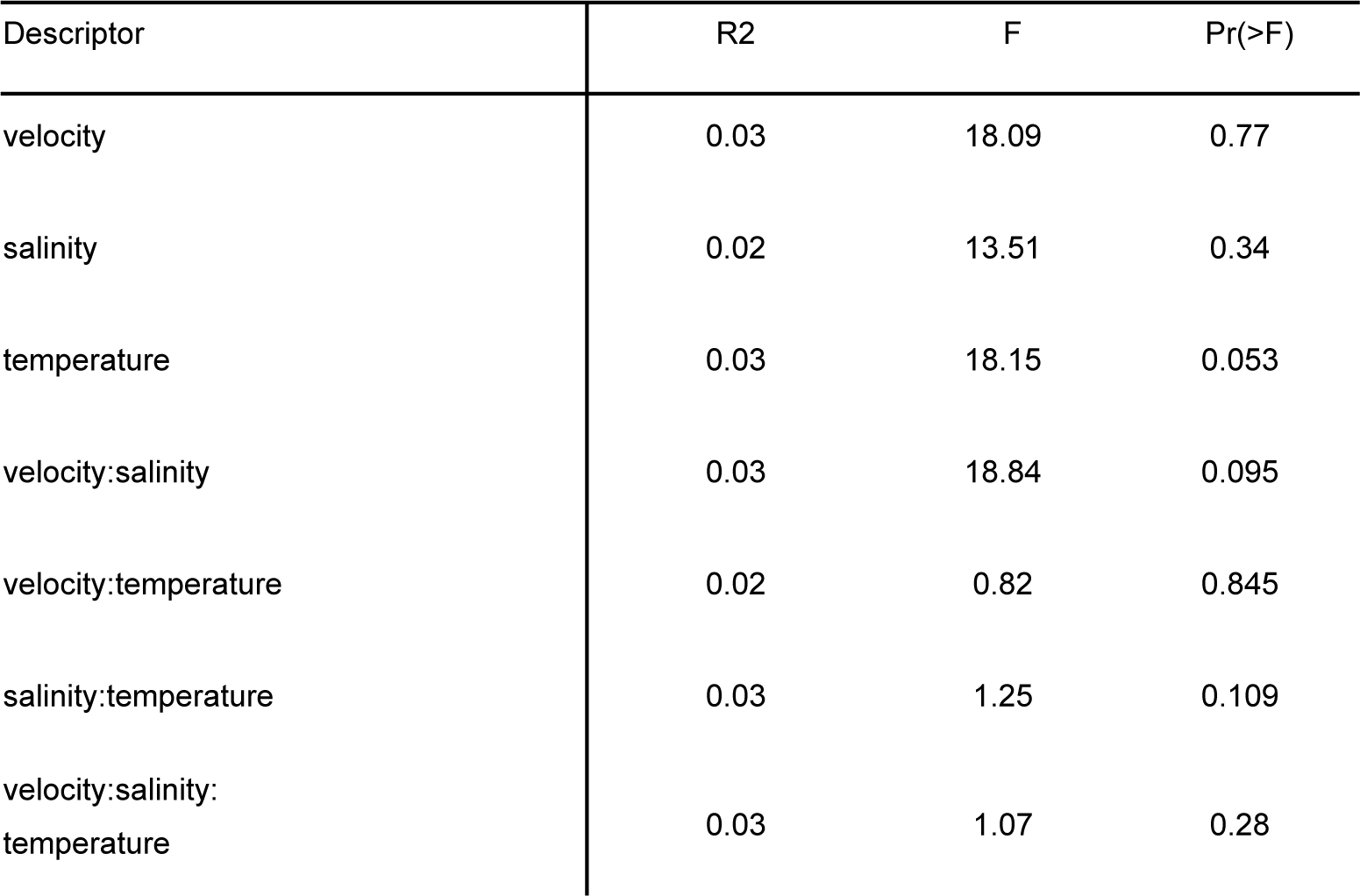
Significance testing of OTUs based on 16S rRNA gene data after the recovery phase with adonis2 (PERMANOVA; n=32). Adonis2 was run with 999 permutations and a model including all single factors and interaction terms based on the marginal effects of the terms as test design. DF=1 for all descriptors.

| Descriptor | R2 | F | Pr(>F) |
| --- | --- | --- | --- |
| velocity | 0.03 | 18.09 | 0.77 |
| salinity | 0.02 | 13.51 | 0.34 |
| temperature | 0.03 | 18.15 | 0.053 |
| velocity:salinity | 0.03 | 18.84 | 0.095 |
| velocity:temperature | 0.02 | 0.82 | 0.845 |
| salinity:temperature | 0.03 | 1.25 | 0.109 |
| velocity:salinity:temperature | 0.03 | 1.07 | 0.28 |

### Alpha diversity of 16S rRNA gene amplicon sequencing data

**Figure S2:**
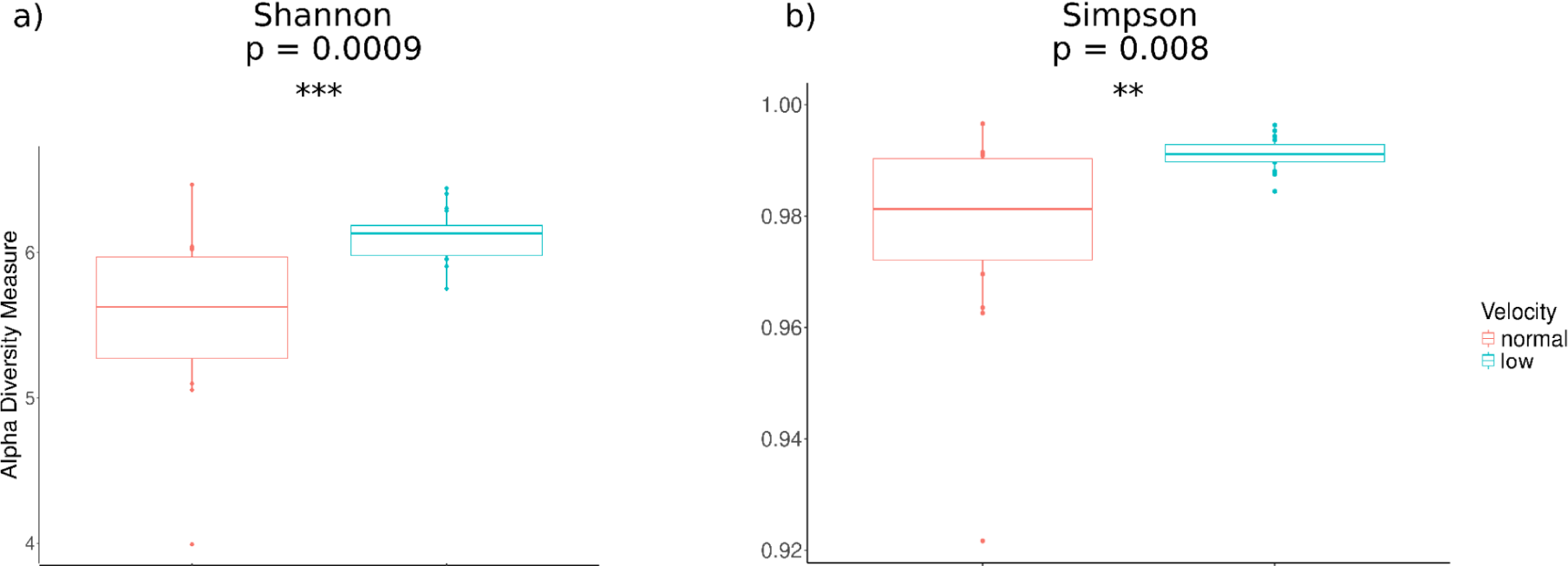
Diversity indices based on bacterial community (16S rRNA amplicon OTUs) under low and normal velocity during the stressor phase. Alpha diversity plot showcases the within-sample diversity of bacterial communities. The x-axis corresponds to samples from lowered and normal velocities, while the y-axis captures alpha diversity indices, offering insights into species richness and evenness. The significance of velocity response was analyzed with GLS estimations.

### Nonpareil curves of metagenomic samples

The diversity of the microbial community and its covered fraction by metagenomic sequencing was estimated using Nonpareil3 (Rodriguez-R et al., 2018).

**Figure S3:**
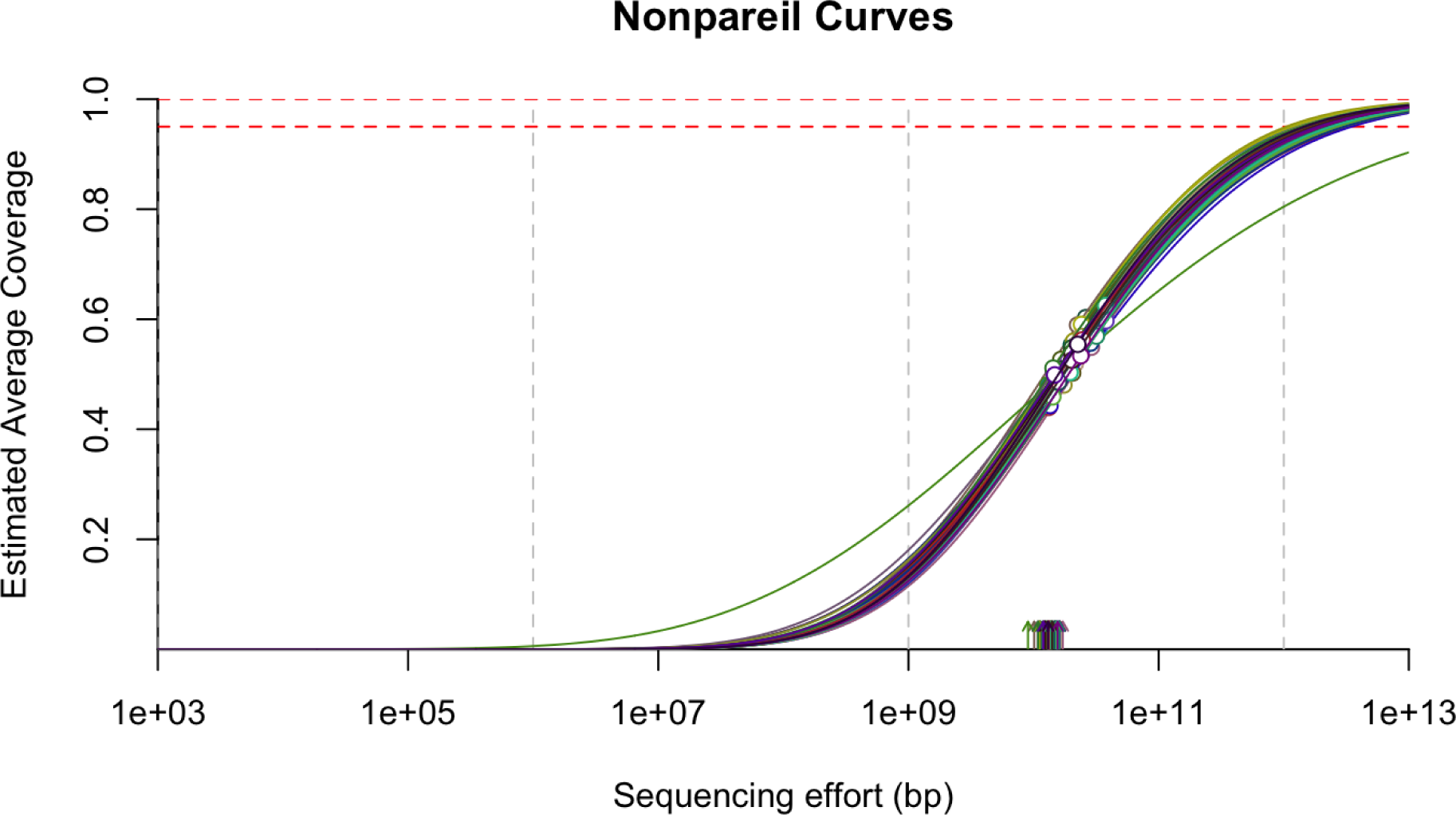
Fitted nonpareil curves for metagenomic samples. Empty circles indicate the actual sequencing effort and arrows the community diversity. Horizontal red dashed lines represent 95% and 99% coverage, respectively. All samples investigated had more than 45% coverage in diversity (average of 54.7 ± 3.77 %). Coverage and diversity estimations by Nonpareil are based on the redundancy of reads in metagenomic datasets; plotted curves represent the fitted model of actual (until dot) and needed sequencing effort to achieve complete coverage of diversity.

### Most abundant microbes across all samples with full taxonomy information

**Figure S4:**
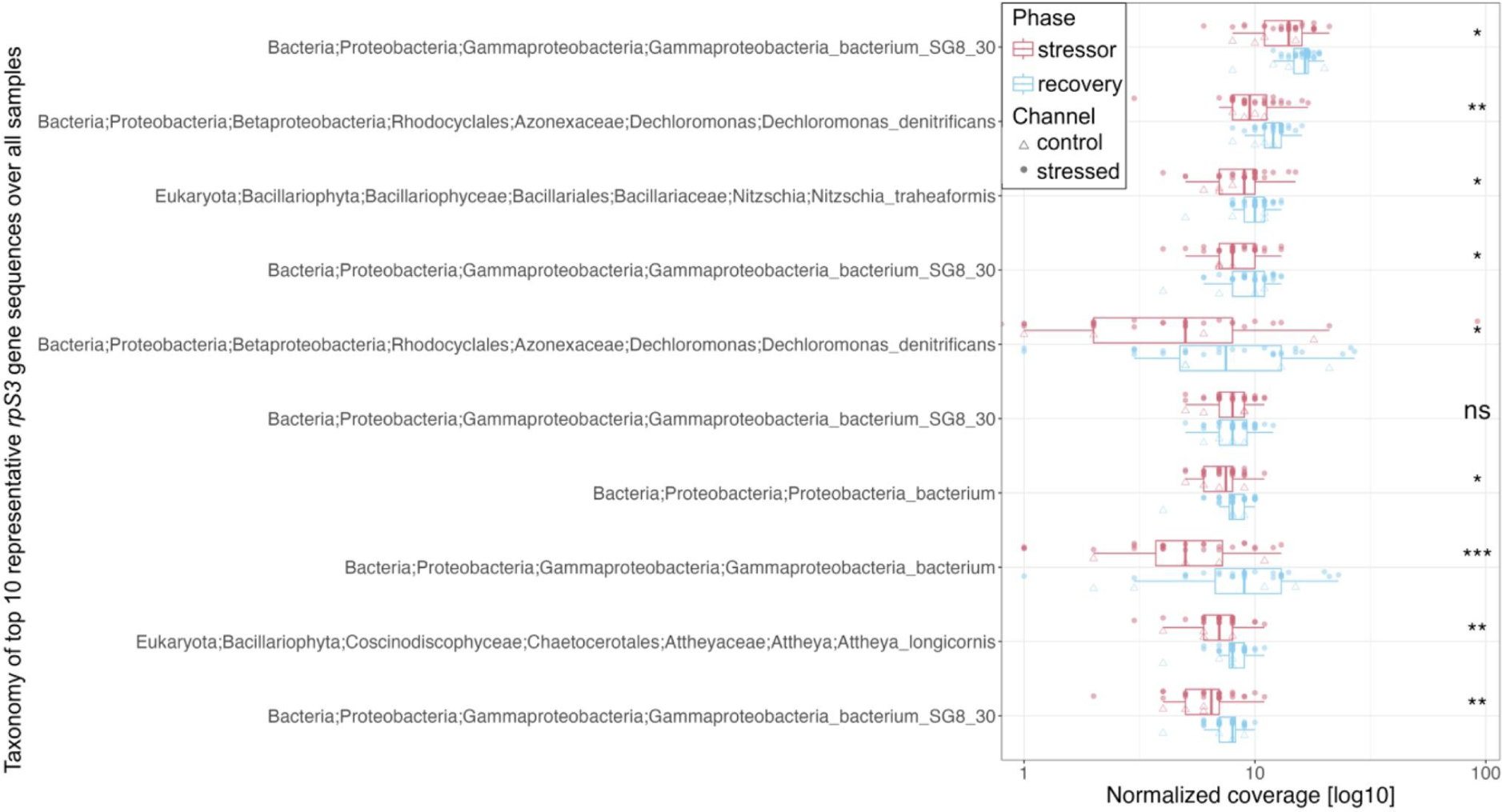
Most abundant microbes across all samples (ordered from top to bottom based on abundance) with full taxonomy information. Sequencing-depth normalized counts of representative *rpS3* gene sequences were summed up over all samples and taxonomy of highest ten genes was annotated. Differences between stressor and recovery phase were tested using the Wilcoxon test.

### Species-specific stressor response based on rpS3 gene analysis

**Figure S5:**
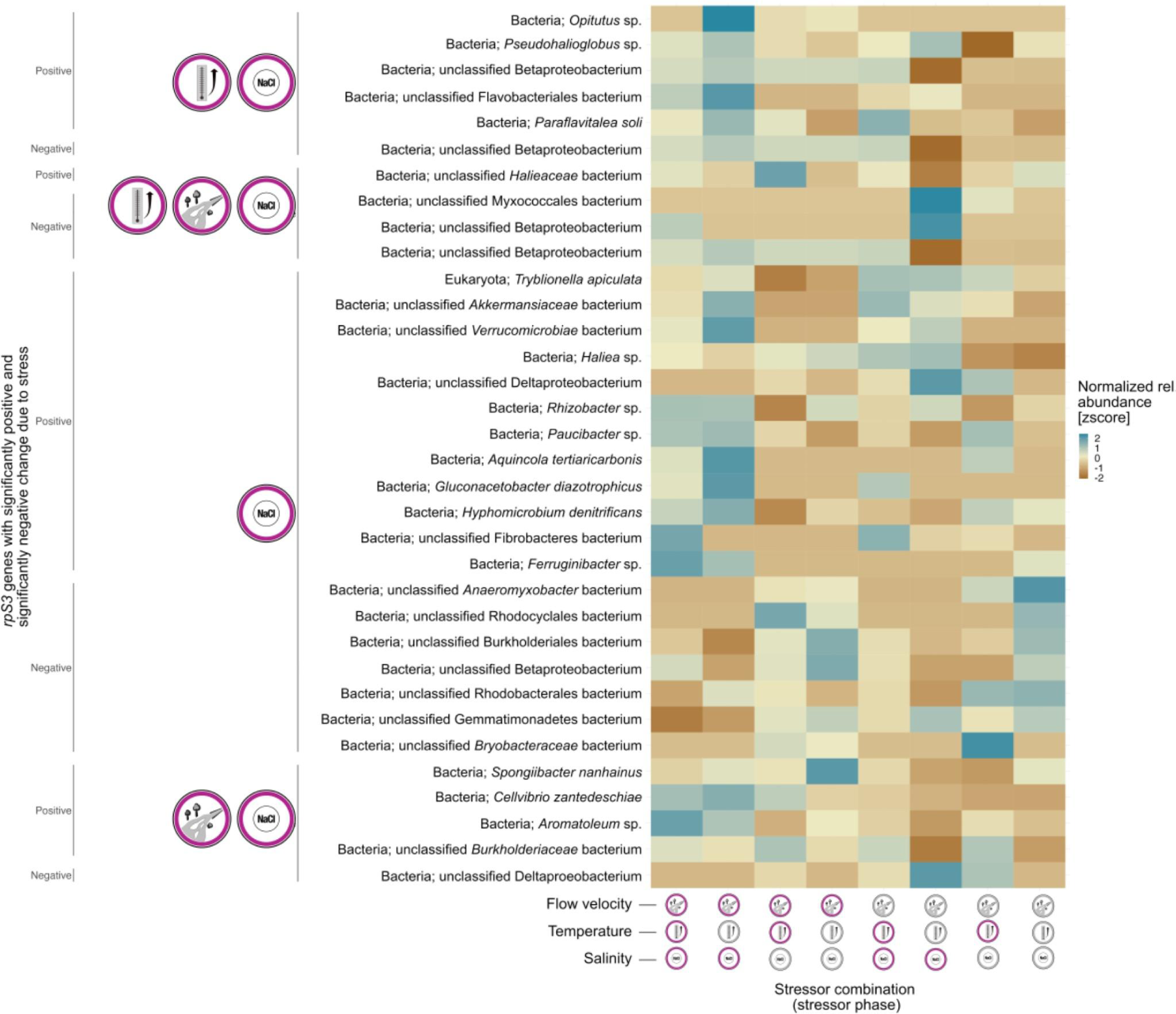
Microbial taxa sensitive to increased salinity based on sequencing-depth normalized representative *rpS3* gene sequences (ANOVA followed by TukeyHSD; adjusted p-value < 0.01). If multiple stressor effects per taxon were significant, only the lowest p-value was chosen.

### Metatranscriptomic analysis of MAGs

**Figure S6:**
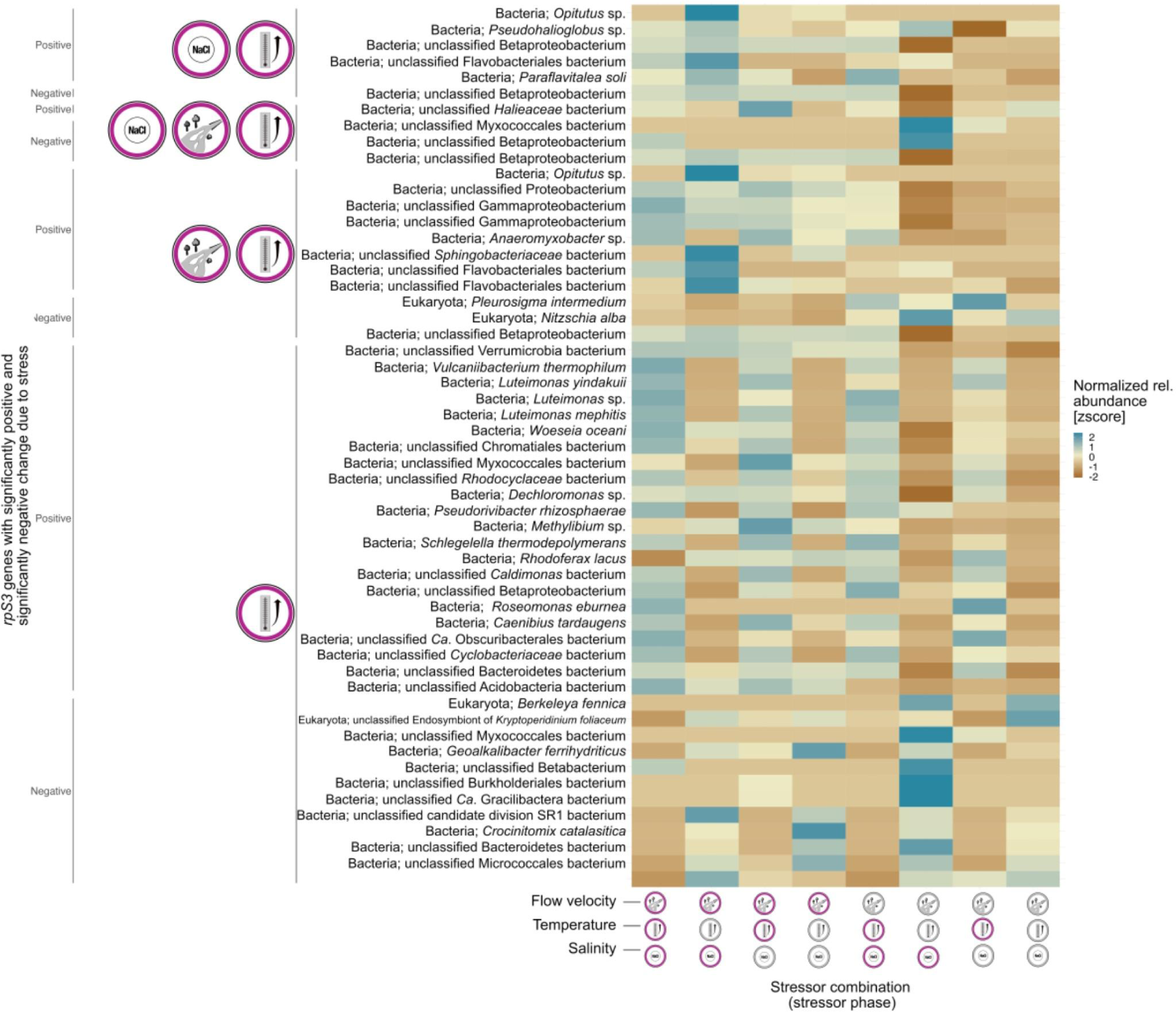
Microbial taxa sensitive to increased temperature based on sequencing-depth normalized representative *rpS3* gene sequences (ANOVA followed by TukeyHSD; adjusted p-value < 0.01). If multiple stressor effects per taxon were significant, only the lowest p-value was chosen.

**Figure S7:**
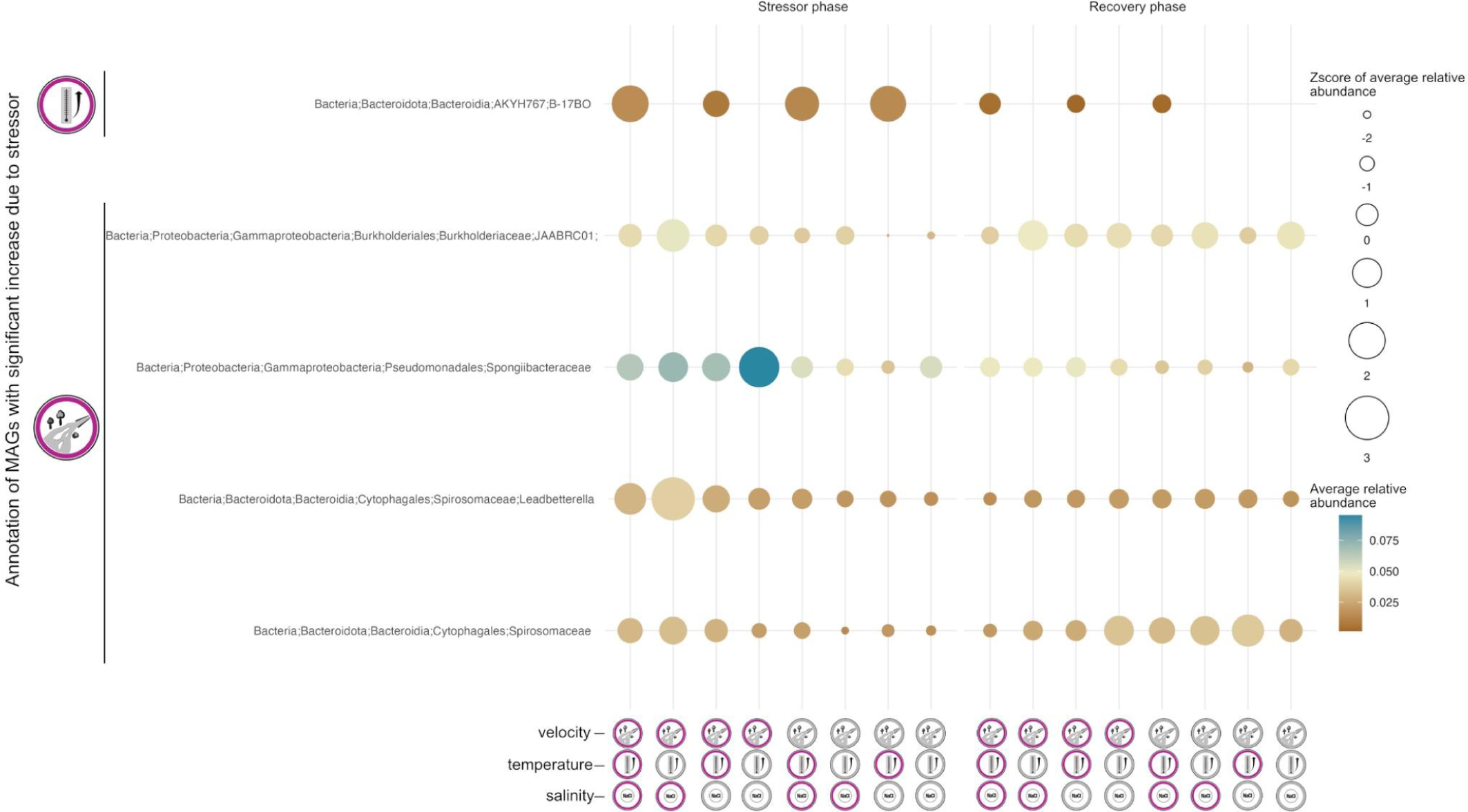
Relative abundance (> 0.1 %) of MAGs with significant response across all mesocosms (n=64), i.e., three MAGs responded positively to lowered flow velocity and one MAG to temperature increase. The latter one was only present in samples with increased temperature indicating a sentinel species.

**Figure S8:**
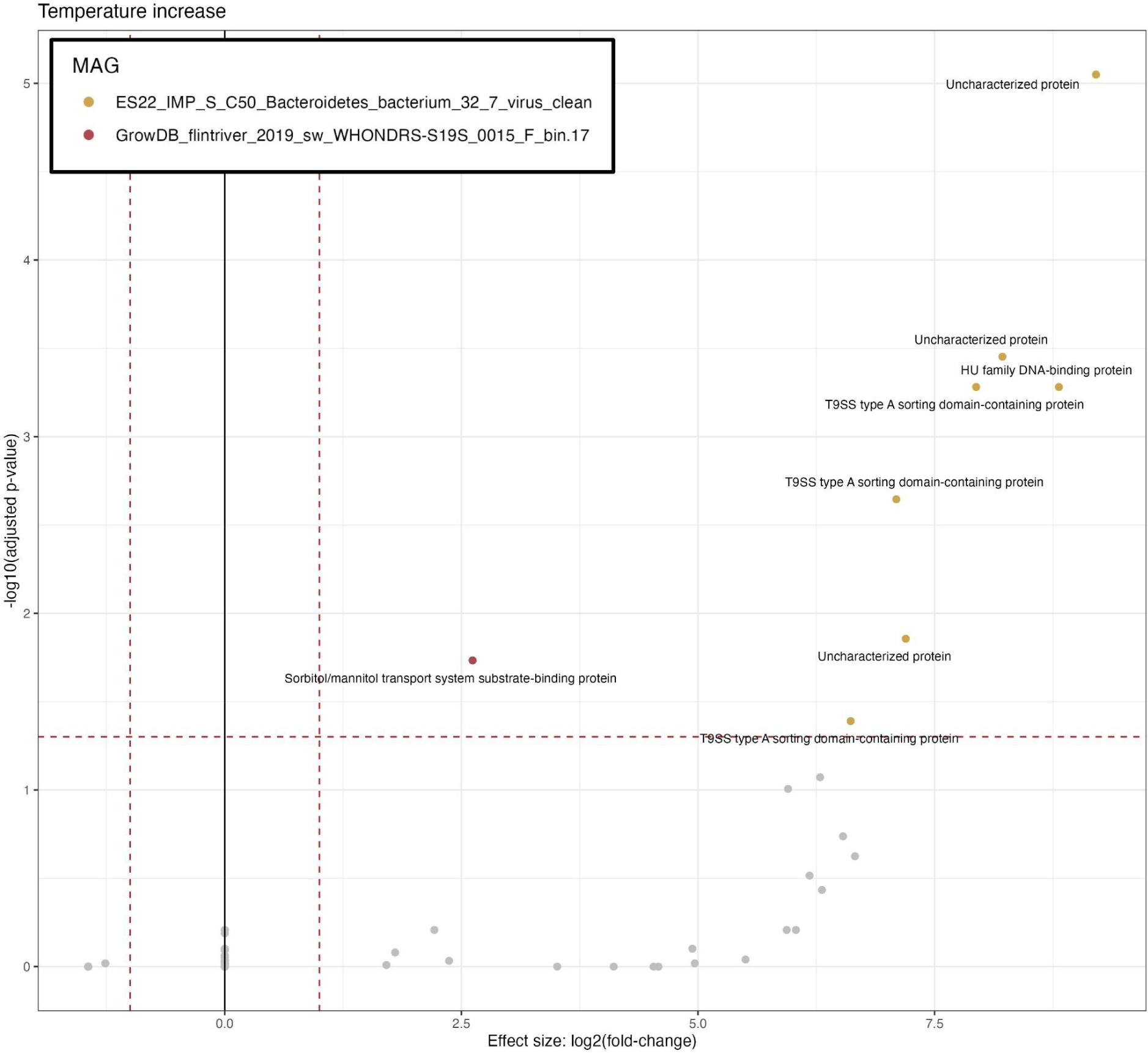
Metatranscriptomic reads were mapped to the set of MAGs and tested for differential expression due to temperature increase using DESeq2 (n=32 for stressor phase; log2(fold-change) > 1 and adjusted p-value < 0.05; Love et al., 2014).

### Statistical testing of encoded ecosystem functions

**Table S2:** Statistical testing of stressor impact on microbial ecosystem functions from metagenomic sequencing. Counts of functional HMMs annotated to metagenomic assemblies by METABOLIC (v4.0) (Zhou et al., 2022) were normalized by sequencing depth. Adonis2 was run with 999 permutations, a model including all single factors and interaction terms with strata set for stressor and recovery phase and based on the marginal effects of the terms as test design.

| Descriptor | Df | SumOfSqs | R2 | F | Pr(>F) |
| --- | --- | --- | --- | --- | --- |
| temperature | 1 | 0.05874 | 0.04 | 2.25 | 0.123 |
| salinity | 1 | 0.03760 | 0.02 | 1.44 | 0.207 |
| velocity | 1 | 0.01730 | 0.01 | 0.66 | 0.444 |
| temperature:salinity | 1 | 0.07567 | 0.05 | 3.05 | 0.067 |
| temperature:velocity | 1 | 0.00628 | 0.004 | 0.25 | 0.709 |
| salinity:velocity | 1 | 0.06602 | 0.04 | 2.66 | 0.091 |
| temperature:salinity:velocity | 1 | 0.05104 | 0.03 | 2.10 | 0.146 |

**Figure S9:**
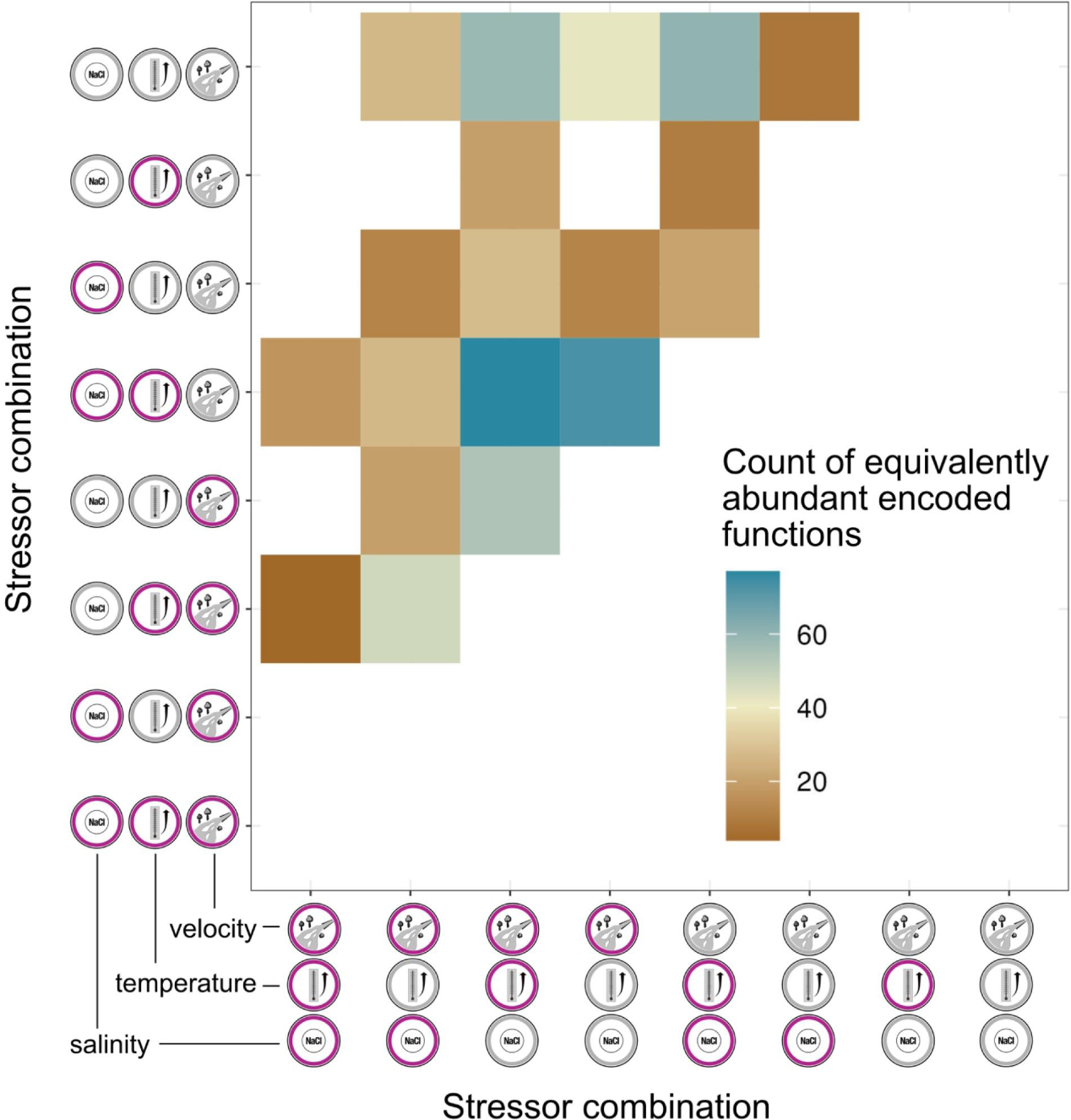
Functional analysis of metagenomic data. Metagenomic assemblies were tested for equivalent abundant functions. Count of significantly equivalently abundant metabolic modules (of in total 314) between the respective stressor pairs (n=32) after the stressor phase indicates that lesser functions are equally shared when more stressors are applied

## Supplementary materials and methods

### Contamination assessment of ExStream mesocosm system

Three single ExStream channels were tested for contamination by filtering circulated flow-through water followed by DNA extraction and Nanopore 16S rRNA gene sequencing. The outflow of an empty channel was connected with a filter system (0.1 mm pore size, JVWP14225) where filtered water was fed again into the mesocosm. This setup was run for nine hours with 10 liter ultrapure water. After cleaning the piping system with diluted bleach, the check was repeated two times with different mesocosm channels. Extraction of DNA was performed based on the DNeasy PowerMax Soil Kit (Qiagen, Germany) with overnight precipitation in 70% EtOH and glycogen as the carrier. Although extracted DNA amounts were too low, 16S rRNA gene sequencing (16S Barcoding Kit 1-24 SQK-16S024, Nanopore) was performed with maximum DNA input resulting in under 30 reads per sample. With working positive and negative controls, no biological contamination was detectable.

### Cycling conditions for 16S rRNA gene amplification

**Table S3:** Cycling conditions for 16S rRNA gene amplification. The first PCR was run for efficient amplification using the target group primers while the second step was performed for individual tagging of the samples and addition of the Illumina adapters. Per sample a reaction volume of 10 µl was used. For 1st PCR: 5 µl Multiplex Master Mix, 0.2 µl forward/reverse primer, 1 µl sample DNA, 3.6 µl water. For 2nd PCR: 5 µl Multiplex Master Mix, 1 µl Coral Load, 1 µl forward/reverse primer, 2 µl sample DNA.

| PCR phase | Time | 1st PCR-Temperature [°C] | 2nd PCR-Temperature [°C] |
| --- | --- | --- | --- |
|  |  | 20 cycles | 25 cycles |
| Initial denaturation | 5 min | 95 | 95 |
| Denaturation (25 cycles) | 30 s | 95 | 95 |
| Annealing (25 cycles) | 90 s | 50 | 61 |
| Elongation (25 cycles) | 30 s | 72 | 72 |
| Final elongation | 10 min | 68 | 68 |

